# Anatomical and molecular organization of the nociceptive medial thalamus

**DOI:** 10.64898/2025.12.05.692629

**Authors:** Lindsay L. Ejoh, Raquel Adaia Sandoval Ortega, Jacqueline W. K. Wu, Alaa Elhiraika, Malaika Mahmood, Kyungdong Kim, Emmy Li, Oliver Joseph, Lisa M. Wooldridge, Jessica A. Wojick, Gregory J. Salimando, Gregory Corder

**Author notes:** Correspondence (G.C.).

## Abstract

The medial thalamus (MTh) is a critical hub for integrating the affective and motivational dimensions of pain, yet its cellular and circuit organization remains poorly defined. Here, we combine activity-dependent genetic tagging, whole-brain clearing and light-sheet imaging, single-cell transcriptomics, and viral circuit tracing to build a comprehensive anatomical and molecular atlas of nociceptive and opioidergic neurons within the MTh. We show that acute and chronic pain activate distinct, spatially localized microdomains across multiple MTh subnuclei, including the central medial, interanteromedial, mediodorsal, and xiphoid nuclei. Activity-dependent tagging reveals a stable ensemble of nociceptive neurons that remains engaged across acute and chronic pain states, suggesting persistent thalamic encoding of affective nociception. Using fluorescent in situ hybridization, we find that most nociceptive-activated neurons express *Oprm1*, and that nearly all *Oprm1*+ neurons are activated by nociceptive stimuli, identifying a pain-active, µ-opioid receptor-expressing population in MTh (MTh^MOR^). Rabies-based tracing demonstrates that MTh^MOR^ neurons receive convergent inputs from nociceptive and inhibitory regions including the anterior cingulate cortex, periaqueductal gray, parabrachial nucleus, and habenula, while projecting broadly to cortical, subcortical, and brainstem structures involved in pain, arousal, and motivation. Enkephalinergic inputs to MTh arise from distinct, non-nociceptive populations, suggesting parallel pathways for nociceptive transmission and endogenous opioid modulation. Together, these findings define the anatomical and molecular organization of the nociceptive medial thalamus and highlight MTh^MOR^ neurons as a cell-type-specific substrate for the development of targeted opioid-based pain therapies.

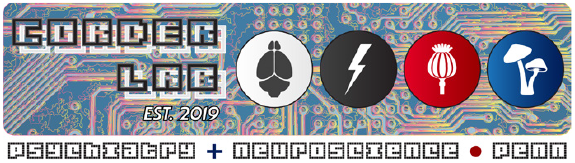

## Introduction

Pain is an aversive experience, resulting from the detection of bodily harm, containing both sensory-discriminative and affective-motivational components^1^. These two components are processed in distinct, brain-wide neural circuits^2,3^, and understanding these discrete features of pain and their underlying circuitry, is crucial for the development of effective analgesics for individuals living with and suffering from chronic pain.

One key brain area receiving this parallel pain information is the thalamus^4^. The spinothalamic tract transmits sensory-discriminative nociceptive information from the spinal cord to the lateral thalamus (e.g., ventral posterolateral, ventral posteromedial, posterior nuclei) for transmission to the primary somatosensory cortex^3,4^, while spinal nociception contributing to the affective-component of pain is partially transmitted through a medial pathway to multiple nuclei of the medial thalamus (MTh)^5,6,7^, before being relayed to various prefrontal cortical areas, amygdala, and dorsal striatum^8,9^. Additionally, MTh nuclei receive nociceptive inputs directly from other brain areas like the parabrachial nucleus^10,11^ and the periaqueductal gray^12,13^

Interestingly, in humans, lesions to lateral thalamic nuclei induces chronic pain in Thalamic Pain Syndrome^14,15^, while lesioning medial thalamus relieves chronic pain symptoms, further suggesting compartmentalization of thalamic nociceptive function^16,17^. Despite this, nociceptive processing in MTh is understudied compared to than lateral thalamic nuclei. The MTh is divided into ten distinct subregions^18,19^ central medial (CM), central lateral (CL), interanteromedial (IAM), intermediodorsal (IMD), mediodorsal (MD) (further broken down the medial (MDm), central (MDc), and lateral (MDl)), paracentral (PC), paraventricular (PV), reuniens (Re), rhomboid (Rh), and xiphoid nucleus (Xi). Numerous studies find increased MTh neural activity to noxious stimuli^4^. For example, a variety of *in vivo* and *ex vivo* recording approaches have observed nociceptive activity in response to electrical shocks^20^, pinches^10,21^, hot temperature^10,22^, cold temperature^23^, visceral pain induced by colorectal distention and chemical irritants like acetic acid^6^, as well as electrical stimulation of peripheral nociceptors^24^. These pain models and recording techniques were independently tested across various single subnuclei, but, to date, none have simultaneously mapped nociceptive neural populations across the entire collection of MTh nuclei.

Importantly, elevated nociceptive activity in MTh nuclei is necessary for affective pain responses in animals and humans. Electric and chemical lesions of various MTh nuclei lead to a reduction in affective behavioral responses to multimodal, visceral and somatic nociceptive stimuli^4,613^, as well as reduction in nociceptive neural activity in cortical areas^24^, illustrating the importance of affective nociceptive transmission to MTh and its recurrent connected areas. Additionally, therapeutic lesions to posterior central lateral and central medial nuclei performed in human patients have modest but significant effects in treating chronic pain^16^. These results highlight the MTh as a key player in the medial nociceptive pathway for both acute and chronic pain with therapeutic potential.

Opioid analgesics, such as morphine and fentanyl, are commonly-prescribed treatments for reducing pain^25^, which carry an array of potential deleterious side-effects, including addiction and respiratory depression ^26^. Recently, we and others have demonstrated that opioids are most efficacious toward the affective component of pain^27–29^, suggesting that functionally-identified, affective nociceptive neurons intersect with molecularly-defined populations expressing the µ-opioid receptor (MOR)^2^. MORs mediate the analgesic effects of exogenous opioids, and endogenous MOR peptide agonists like endorphins and enkephalins^30,31^. Additionally, nociceptive activity in MTh diminishes during endogenous forms of pain relief and stress-induced analgesia and placebo analgesia attenuate MTh activity in a MOR-dependent manner in both human^32–34^ and animal models^35^. However, it remains unknown from which brain regions endogenous opioids ligands may be released into MTh. This study aims to characterize nociceptive responses in MTh substructures, quantify and compare afferents to and efferents from MOR^+^ medial thalamic cells (MTh-^MOR^), with a focus on cell-type specific afferents that may mediate endogenous analgesia.

The MTh expresses one of the highest densities of MORs in the brain^36,37^. Opioid agonists reduce activity in MTh^MOR^ neurons, shifting from a tonic to burst firing mode, a pattern often associated with analgesia^38,39^. Microinjection of opioids into MTh also relieves affective-motivational pain behaviors^20^. These results indicate that MTh^MOR^ neurons might be a useful cell type for mediating pain relief. However, the responses of MTh^MOR^ cells during nociception remains vastly under-explored. Targeting MOR-expressing cell types (MOR^+^) in specific brain circuits for affective pain processes may open an avenue to a potential new class of analgesics that diminish the unpleasantness associated with pain rather than pain sensation without the associated addictive and respiratory depressive side effects of opioid drugs^40,41,42^.

Due to the lack of an investigation of all MTh nuclei simultaneously during pain, it remains unclear if there are anatomically-restricted ‘pain hotspots’ in MTh, in which there is preferential activation of certain MTh nuclei in response to pain, or if MTh nuclei respond uniformly to nociceptive stimuli. It also is unknown if MTh nociceptive responses differ with acute and chronic pain, whether it is the pain-responsive MTh neurons that express MOR, and whether previously-discovered inputs to and outputs from MTh are maintained in the MTh^MOR^ population.

In the present study, we combine activity-dependent immediate early gene mapping, transcriptomics, and rabies and axonal circuit tracing tools to uncover the nature of opioidergic circuitry in the nociceptive MTh. We show that the medial thalamus contains a distinct and stable population of pain-modulated, MOR^+^ neurons that receive converging inputs from cortical and subcortical regions. The present study reveals novel information about how pain is processed and potentially modulated in the thalamus, as well as a novel circuit to target for the development of personalized therapies to mimic analgesia without the adverse side effects found with traditional opioid pharmacotherapies.

## Results

### Acute nociception and allodynia increase MTh activity

To quantitatively map nociceptive neuronal engagement across the ten MTh subnuclei during acute and chronic pain states, we leveraged lipid-cleared brain imaging to localize expression of the noxious-evoked, activity-dependent immediate early gene, FOS, across the 3-dimensional volume of MTh structures. First, we used TRAP2 mice^43^ (*Fos*-FOS-2A-iCre^ERT2^) crossed with a nuclear Sun1-sfGFP fluorescent reporter line to permanently label MTh neuronal ensembles activated by noxious hot water stimuli (acute painTRAP; **Fig. 1A**). For acute painTRAP, we applied repeated 55°C water droplets to the left hind paw (20 drops (∼50 μl per drop), once every 30 seconds for 10 minutes). 60-min post-stimuli, we administered 4-hydroxytamoxifen (4-OHT; 40 mg/kg subcutaneous) to initiate Cre-recombination to induce permanent expression of nuclear sfGFP in FOS-expressing MTh neurons. Three days after acute painTRAP, mice were either injured via transection of the left sciatic nerve (SNI, Spared Nerve Injury model) to induce chronic neuropathic pain or remained uninjured

**Fig. 1.**
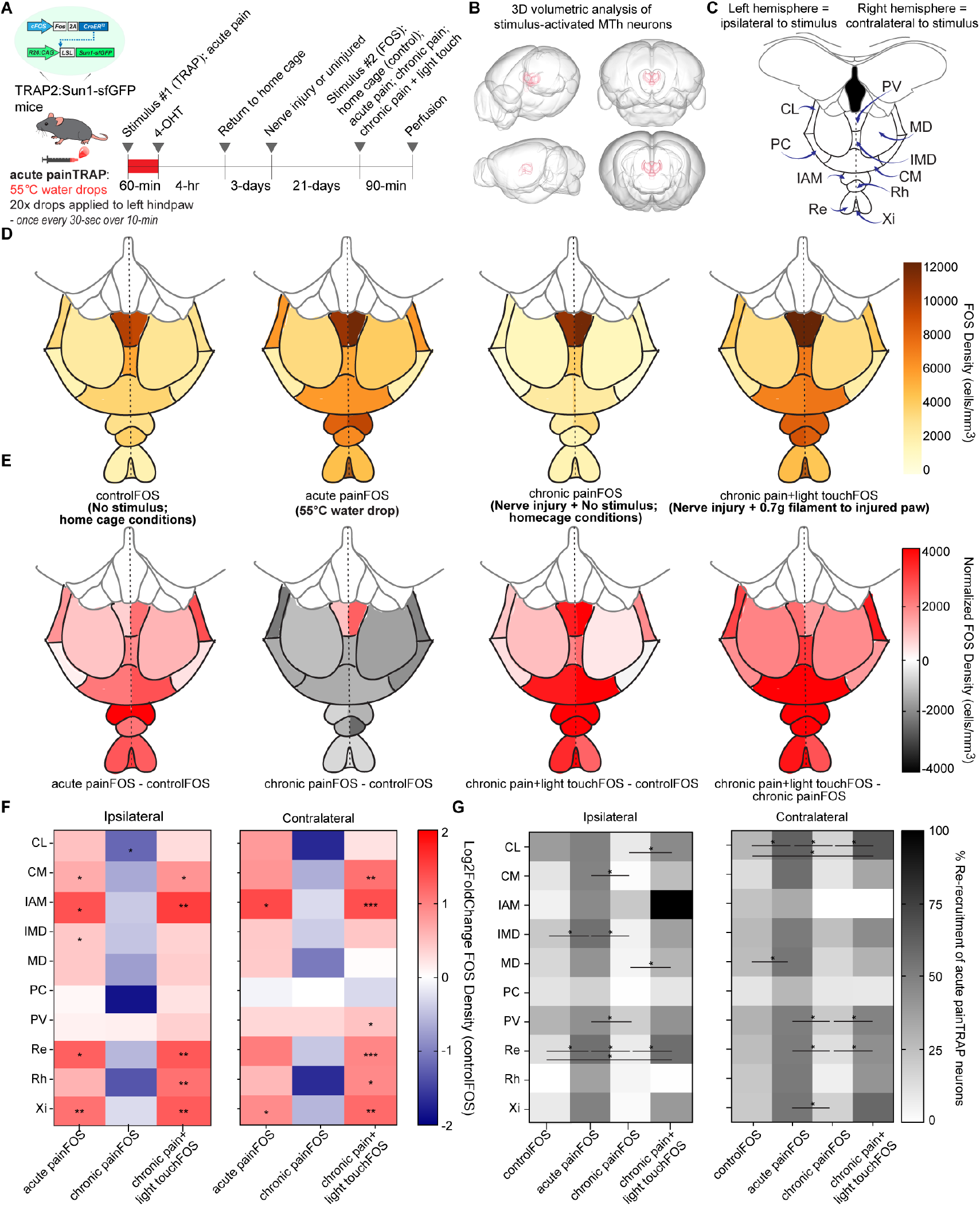
Acute nociception and allodynia increase MTh activity. (A) Experimental design for whole-brain clearing and light sheet imaging of acute painTRAP^sfGFP^ labeled nuclei, as well as FOS-labeled nuclei with either an acute nociceptive stimulus (acute painFOS; n=4 males), allodynia during chronic neuropathic pain (chronic pain + light touchFOS; n=4 males), or home cage groups during injured (chronic painFOS; n=3 males) or uninjured (controlFOS; n=4 males) states. (B) 3D renderings of medial thalamic nuclei obtained from Howard Hughes Medical Institute’s Janelia’s MouseLight Neuron Browser. (C) Coronal cross-section atlas of medial thalamus annotated with the ten MTh subnuclei, each split into ipsilateral (same hemisphere as stimulus) and contralateral (opposite hemisphere as stimulus) parts. (D) Spatial heat maps of mean density of FOS-expressing cells in MTh in each group. (G) Heatmap for mean percent of acute painTRAP^sf-GFP^ co-labeled with FOS for each group, calculated using: (number of co-labeled cells / number of painTRAP^sf-GFP^ cells)*100%. Two-way ANOVA with Bonferonni multiple comparisons test determined significance *p<0.05. Abbreviations: CL, central lateral. CM, central medial nucleus. IAM, interanteromedial nucleus. IMD, intermediodorsal nucleus. MD, mediodorsal thalamus PC, paracentral nucleus. PV, paraventricular nucleus. Re, nucleus reuniens. Rh, rhomboid nucleus. Xi, xiphoid nucleus.

After a 21-day period to allow the development of chronic neuropathic pain, mice were randomly assigned to one of four experimental groups (**Fig. S1A**) and exposed to a second stimulus: 1) controlFOS – uninjured mice that remained in their home cages without hindpaw stimulation; 2) acute painFOS – uninjured mice that received repeated noxious 55 °C water droplets applied to the left hindpaw; 3) chronic painFOS – SNI mice that remained in their home cages without stimulation; and 4) chronic pain + light touch FOS – SNI mice that received a light tactile stimulus (0.16 g von Frey filament, 20 applications delivered once every 30 s for 10 min) to the hypersensitive hindpaw to evoke mechanical allodynia.

Ninety minutes after stimulus delivery—allowing sufficient time for induction of FOS^44^—brains were collected and processed for SHIELD lipid clearing, immunohistochemistry for anti-GFP and anti-FOS, and volumetric light-sheet imaging. This enabled visualization of sfGFP+ nuclei (neurons activated by the initial acute painTRAP stimulus three weeks earlier) and FOS+ nuclei (neurons activated by the second stimulus) across both hemispheres of the MTh in three dimensions (**Fig. 1B–C**). Cleared brains were registered to the Allen CCFv3 atlas for automated quantification. Spa-tial heat map analysis illustrates an overview of FOS densities between control and nociceptive stimuli **(Fig. 1D)**, as well as FOS densities when normalized to home cage control conditions **(Fig. 1E)**.

We found that the acute nociceptive stimulus increased FOS expression in several MTh subnuclei, across hemispheres ipsilateral and contralateral to the left hind paw stimuli **(Fig. 1F, S1C, S2A)**. For example, relative to controlFOS mice, acute painFOS mice showed a significant ∼3-fold higher density of FOS^+^ cells in the contralateral IAM **(Fig. S2A)**. We then normalized FOS density by calculating the log2fold change of density of FOS^+^ cells in the acute painFOS group compared to uninjured controlFOS mice. After normalization, we identified increased nociceptive activity in the contralateral IAM and Xi **(Fig. 1F; S2E)**, and ipsilateral CM, IAM, IMD, Re, Xi nuclei **(Fig. 1F, S1G)**. These data demonstrate that MTh nuclei do not respond uniformly to acute nociceptive stimuli.

After investigating responses to acute nociception in MTh, we used the chronic pain and chronic pain + light touch groups to assess activity differences during chronic nociception. First, to confirm whether activity in MTh is altered during a resting chronic pain state, we compared FOS density between the chronic pain and controlFOS conditions. Interestingly, though there were trends towards a reduction in FOS density after SNI, we did not observe any MTh nuclei with significant reduction in FOS density when observing pair-wise differences between the neuropathic pain and uninjured home cage conditions **(S1D, S2B)**. However, normalization to uninjured controlFOS conditions revealed a significant reduction in activity in the CL ipsilateral to the nociceptive stimulus **(Fig. 1F; Fig. S1G)**.

Neuropathic pain results in symptoms of mechanical allodynia, in which innocuous stimuli such as gentle, light touch are perceived as noxious^45^. In the chronic pain + light touchFOS condition, we found elevated FOS compared to controlFOS mice, with significant increases in contralateral CM, IAM, Rh **(Fig. S2C)**, and IAM, Rh, Xi ipsilateral to the light touch stimulus **(Fig. S1E)**. After normalizing to the uninjured home cage condition, we detected significant activation of contralateral CM, IAM, PV, Re, Rh, Xi **(Fig. 1F; S2E)**, and ipsilateral CM, IAM, Re, Rh, Xi subnuclei **(Fig. 1F; S1G)**. Furthermore, we observed elevated chronic pain + light touch FOS density compared to the neuropathic unstimulated chronic pain condition (chronic painFOS) in contralateral CM, IAM, Rh, and Xi **(Fig. S2D)** and ipsilateral CM, IAM, Re, Rh, and Xi **(Fig. S1F)**. Additionally there were no differences in FOS density between the ipsilateral and contralateral hemispheres in any condition **(Fig. S1B)**, indicating the absence of laterality in stimulus-evoked nociceptive responses in MTh.

### Stable nociceptive MTh subnuclei: acute to chronic pain transition states

Understanding the stability of neural activity over days is crucial for addressing the concept of “representational drift”—the idea that the neural representation of sensory or behavioral information changes over time^46^. Determining whether the nociceptive neurons within each MTh subnuclei maintains stable nociceptive activity across the acute to chronic pain transition period may be key for developing pharmacological pain therapies targeting specific functional or molecular pathways in this ensemble^47,48^. First, to assess if acute painTRAP cells are re-engaged by subsequent bouts of identical acute nociceptive stimuli, we quantified the proportion of acute painTRAP^sf-GFP+^ neurons that co-expressed acute painFOS. Compared to controlFOS conditions, we found significantly more acute noxiously-re-engaged neurons in ipsilateral IMD and Re **(Fig. 1G; S1H)**, and importantlycontralateral CL and MD **(Fig. S1G; S2F)**, suggesting the existence of stable, acute nociceptive populations within these distinct subnuclei.

Next, we sought to determine whether acute nociceptive-active neurons in MTh were re-engaged weeks later following the induction of chronic neuropathic pain. Notably, light touch-evoked re-engagement of the acute painTRAP neurons was significantly increased in the contralateral CL **(Fig. 1G; S2F)**, and ipsilateral Re **(Fig. 1G; S1H)**, indicating the existence of MTh ensembles that activate in response to both acute and chronic nociception.

Our activity-dependent genetic labeling study reveals differential responses to acute pain and neuropathic allodynia among different MTh. However, measuring activity summed throughout the 3D volume of each subnucleus might miss possible nociceptive “hotspots” within specific anatomic coordinates. Therefore, we next investigated whether the MTh nuclei also contain nociceptive hotspots within their anterior-posterior spatial axes.

### Acute and chronic nociception engages cells across the spatial axis of MTh

To investigate the presence of nociceptive hotspots within the spatial axes of MTh subnuclei, we utilized the TRAP2 mouse crossed with an Ai9 tdTomato fluorescent reporter to create the TRAP2:Ai9 mouse that allows for genetic capturing and permanent labeling of nociceptive-activated neural ensembles **(Fig. 2A)**. Animals were divided into two groups: those that received a noxious hot water (55°C) stimulus to label acute nociceptive-activated cells (acute painTRAP) and control mice that remained in a home cage (controlTRAP) **(Fig. 2F)**. Seven days later, animals were either given SNI to the left sciatic nerve or remained uninjured. After 21 days, all mice were exposed to a second stimulus to drive expression of FOS for immunolabeling to map overlap with the original TRAP^+^ tdTomato neurons.

**Fig. 2.**
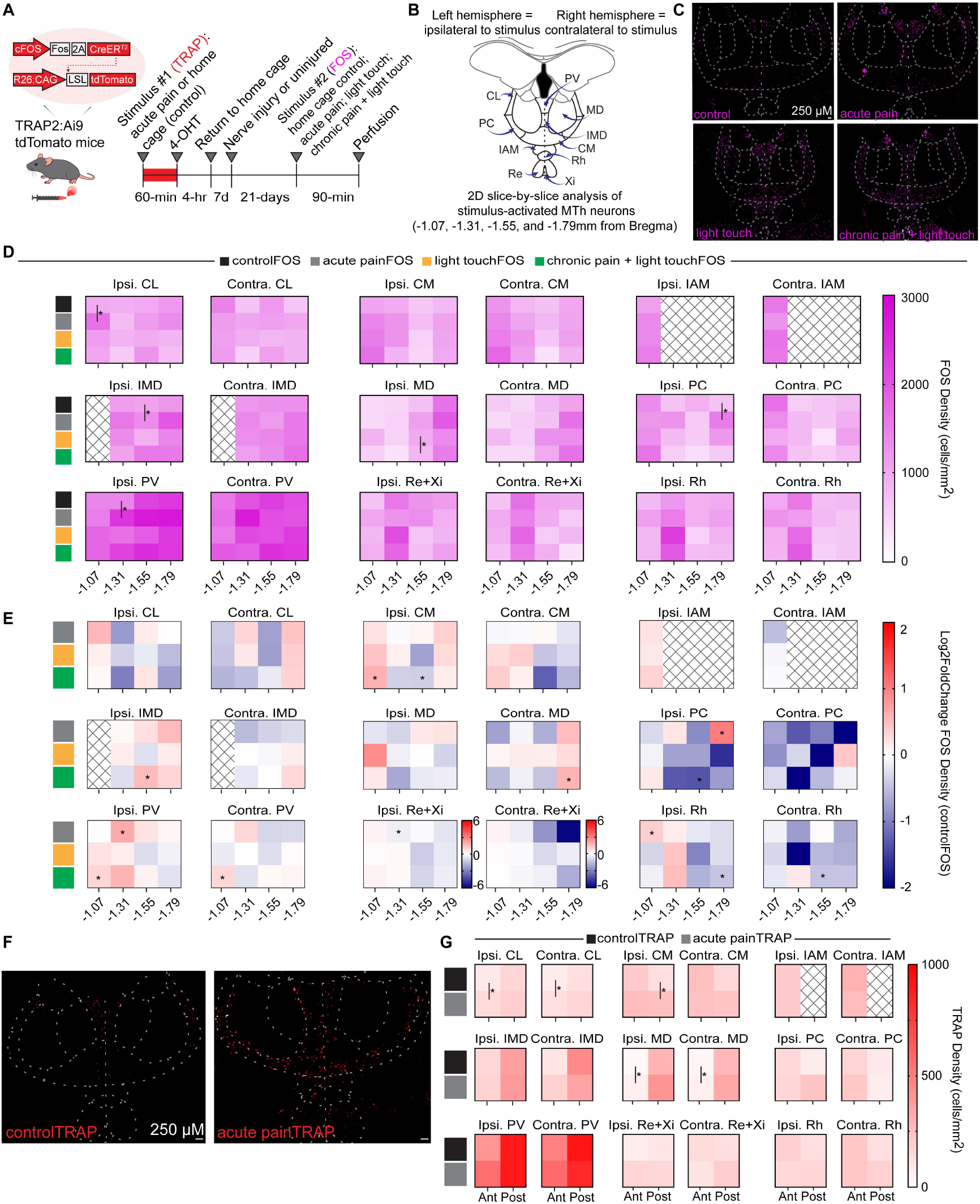
Non-uniform responses to acute and chronic nociception within spatial axes of MTh subnuclei. (A) Experimental design for activity-dependent labeling and quantification of acute painTRAP^Ai9^ (n=11 females, n=5 males) or controlTRAP^Ai9^ (n=11 females, n=4 males)labeled nuclei, as well as FOS-labeled nuclei with either an acute nociceptive stimulus (acute painFOS; n=5 females, n=3 males), allodynia during chronic neuropathic pain (chronic pain + light touchFOS; n=7 females, n=1 male), light touch in an uninjured state (light touchFOS; n=5 females, n=3 males) or home cage groups during an uninjured state (controlFOS; n=5 females, n=2 males). (B) Coronal cross-section atlas of medial thalamus annotated with the ten MTh subnuclei, each split into ipsilateral (same hemisphere as stimulus) and contralateral (opposite hemisphere as stimulus) parts. (C) 20x representative fluorescent images of FOS expression in each group. (D) Heat maps for FOS density in each group at −1.07, −1.31, −1.55, −1.79mm from Bregma in each MTh subnucleus. Independent t-tests between controlFOS and acute painFOS, and light touchFOS and chronic pain + lightouchFOS determined significance *p<0.05. (E) Heat maps for mean Log2 fold change in FOS density for each group at −1.07, −1.31, −1.55, −1.79mm from Bregma in each MTh subnucleus compared to the controlFOS condition; one sample t-test vs. zero determined significance *p<0.05. (F) 20X representative fluorescent images of controlTRAP^Ai9^ and acute painTRAP^Ai9^ in MTh. (G) Heat maps for controlTRAP vs. acute painTRAP density at anterior (−1.07mm and −1.31mm averaged) and posterior (−1.55 and −1.79 averaged) coordinates. Independent t-tests determined significance *p<0.05. (Scale bars: 250 µM). Abbreviations: CL, central lateral. CM, central medial nucleus. IAM, interanteromedial nucleus. IMD, intermediodorsal nucleus. MD, mediodorsal thalamus PC, paracentral nucleus. PV, paraventricular nucleus. Re+Xi, averaged across nucleus reuniens and xiphoid nucleus. Rh, rhomboid nucleus.

Mice were randomly assigned to one of four FOS groups: (1) uninjured mice that remained in their home cages (controlFOS), (2) uninjured mice that received hot water drops to the left hind paw (acute painFOS), (3) uninjured mice that received light touch stimuli to the left hind paw (light touchFOS), and (4) nerve-injured mice that received light touch stimuli to their hypersensitive left hind paw (chronic pain + light touchFOS). Ninety minutes after the FOS-evoking stimulus, mice were perfused, and immunohistochemistry was performed to label tdTomato and FOS across four anterior–posterior levels of the medial thalamus (−1.07 mm, −1.31 mm, −1.55 mm, and −1.79 mm from Bregma; **Fig. 2B-C**).

Acute painFOS engaged significantly more neurons in CL (− 1.07mm), PC (−1.79mm), and PV (−1.31mm) ipsilateral to the stimulus compared to home cage controlFOS conditions **(Fig. 2D)**. After normalizing FOS density to controlFOS conditions, we also identified increased acute nociceptive activity in ipsilateral PC (− 1.79mm), PV (−1.31mm), and Rh (−1.07mm) subnuclei **(Fig. 2E)**. Notably, one area-ipsilateral Re+Xi (−1.31mm)- engaged a significantly lower density of acute painFOS neurons compared to controlFOS **(Fig. 2E)**. We also quantified density of TRAP^Ai9+^ cells across coordinates in MTh nuclei and found no significant differences between controlTRAP and acute painTRAP densities in any coordinate **(Fig. S3)**. However, when combining −1.07mm and −1.31mm into an “anterior” category, and −1.55mm and −1.79mm into a “posterior” category, we observed elevated densities of acute painTRAP^Ai9+^ neurons in both hemispheres of the anterior CL, both hemispheres of the anterior MD, and contralateral anterior CM **(Fig. 2G)**. Notably, both TRAP and FOS-based measures of neural activity reveal that acute nociception engaged more neurons than control conditions in CL thalamus **(Fig. 2D, G)**.

Next, chronic pain + light touchFOS conditions were compared to light touchFOS and controlFOS to investigate the impact of chronic nociception on activity within spatial axes of MTh. Compared to light touchFOS conditions, elevated chronic pain + light touchFOS density was found in ipsilateral MD (−1.55mm; **Fig. 2D**). When normalized to controlFOS conditions, chronic pain + light touch significantly increased activity in contralateral MD (− 1.79mm) and PV (−1.07mm), and ipsilateral CM (−1.07mm), IMD (− 1.55mm), and PV (−1.07mm; **Fig. 2E**). Interestingly, we also observed that chronic pain + light touch reduced activity in posterior coordinates of several nuclei, including contralateral Rh (− 1.55mm) and ipsilateral CM (−1.55mm), PC (−1.55mm), and Rh (− 1.79mm; **Fig. 2E**). Consistent with our 3D analysis, these results demonstrate bilateral engagement of MTh nuclei during both acute and chronic nociception. However, quantification across anterior–posterior coordinates revealed anatomical differences among several nuclei that may underlie the overall 3D effects, uncovered spatially localized patterns not evident in whole-nucleus analyses, and even identified microdomains within MTh subnuclei exhibiting opposing pain-related activity patterns.

### Transcriptomic analysis of MTh reveals µ-opioid receptor expression in acute nociceptive-activated cells

MOR is expressed densely in the MTh **(Fig. 3C)**^36,49^, and µ-opioid agonists-mediated inhibition of MTh neurons is anti-nociceptive, impairing affective nociceptive processing^20^. However, the distribution of MOR^+^ cells throughout MTh subnuclei, and the degree of activation of this specific cell-type by nociceptive stimuli remain unexplored. To address this, we performed fluorescent in situ hybridization (FISH) to visualize acute painTRAP^Ai9+^ and *Oprm1*^+^ mRNA within the MTh (**Fig. 3A–B, D**). Using the HALO deep-learning semi-automated quantification platform, which resolves thousands of individual mRNA transcripts within single cells across large tissue volumes, we mapped the distribution of *Oprm1* mRNA transcript counts across all MTh cells (**Fig. 3E**). We found that approximately 50% of *Oprm1*^*+*^ cells expressed ≥14 transcripts, 25% expressed fewer than 8, and 25% expressed more than 22 transcripts. Based on these differences, cells were classified into four categories: low-expressing (1–8 transcripts), medium-expressing (9–14), high-expressing (15–21), and very high-expressing (≥22). Each MTh subnucleus exhibited a distinct *Oprm1* transcript distribution profile (**Fig. 3F; S5A–L**). Median transcript levels per subnucleus ranged from 12–16, with the central mediodorsal (MDc) and medial mediodorsal (MDm) nuclei showing the highest median counts (16), and the interanteromedial (IAM) and xiphoid (Xi) nuclei showing the lowest (12).

**Fig. 3.**
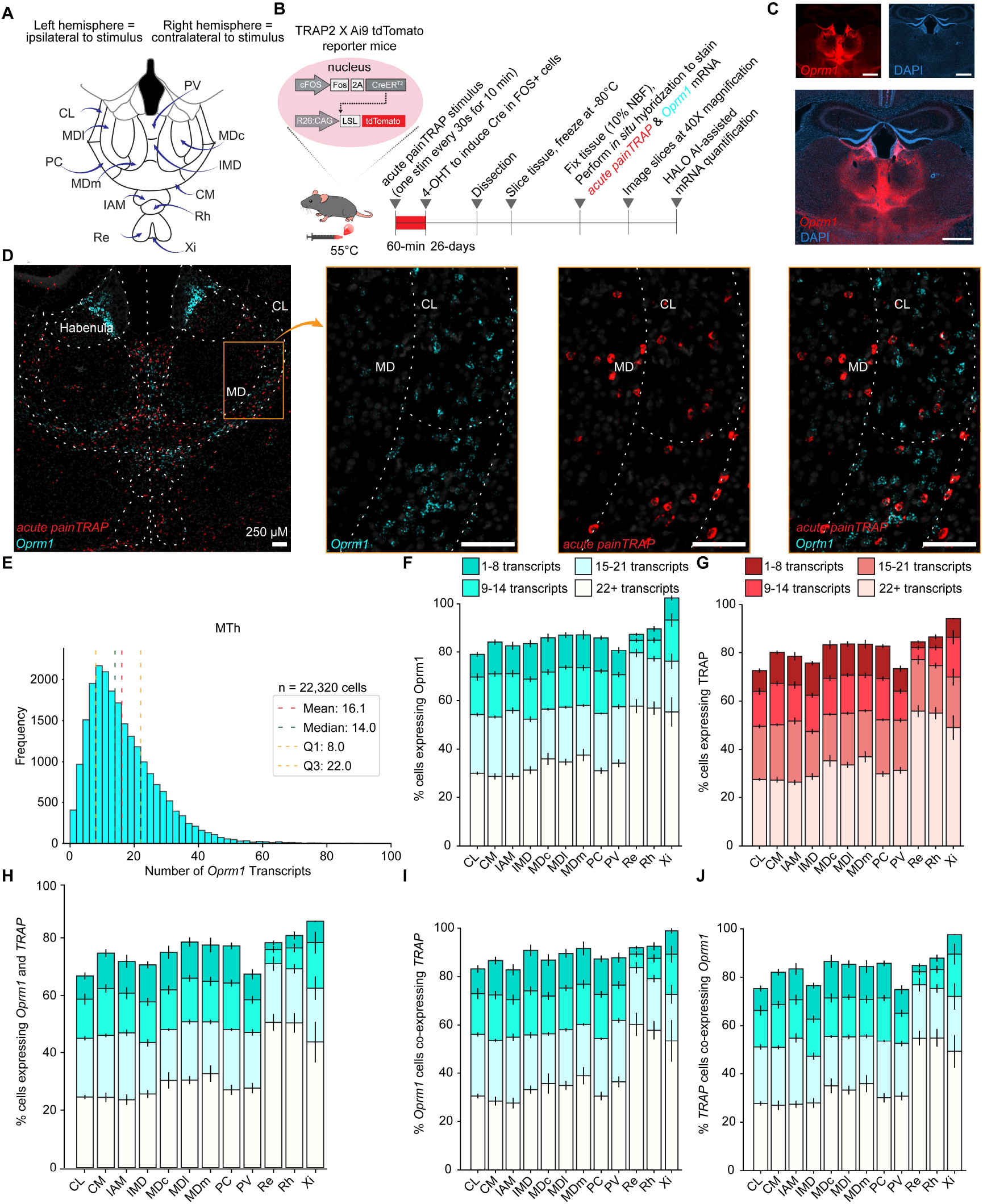
Transcriptomic analysis of MTh reveals nociceptive-activated cells express MOR. (A) Coronal cross-section atlas of medial thalamus annotated with nine MTh subnuclei and the three subnuclei of MD thalamus, each split into ipsilateral (same hemisphere as stimulus) and contralateral (opposite hemisphere as stimulus) parts. (B) TRAP2:Ai9 mice for activity-dependent genetic capturing and permanent tdTomato fluorescent tagging of acute nociceptive cells (*acute painTRAP*^*Ai9+*^). Experimental design for *acute painTRAP* and *in situ* hybridization to label and quantify *acute painTRAP*^*Ai9+*^ and *Oprm1* mRNA in MTh. (C) 20x representative fluorescent images of MOR protein in MTh. (D) 20x representative fluorescent images of mRNA transcripts for *acute painTRAP*^*Ai9*^ (red) and *Oprm1* (light blue) in MTh at approximately A/P −1.31mm relative to Bregma. (E) Histogram for Number of *Oprm1* transcripts throughout entire MTh for n=3 female and n=3 male TRAP2:Ai9 mice. (F) Quantification of the percent of cells expressing *Oprm1* in each MTh subnucleus (Number of *Oprm1*^+^ cells/total number of cells)*100%. (G) Quantification of the percent of cells expressing TRAP in each MTh subnucleus (Number of *acute painTRAP*^*Ai9+*^ cells/total number of cells)*100%. (H) Quantification of the percent of cells expressing both *Oprm1* and *acute painTRAP*^*Ai9*^ in each MTh subnucleus (Number of *acute painTRAP*^*Ai9*^ and *Oprm1* co-expressing cells/total number of cells)*100%. (I) Quantification of the percent of *Oprm1*^+^ cells co-expressing *acute painTRAP*^*Ai9*^ (Number of *acute painTRAP*^*Ai9*^ and *Oprm1* co-expressing cells/total number of *Oprm1*^+^ cells)*100%. (J) Quantification of the percent of *acute painTRAP*^*Ai9+*^ cells co-expressing *Oprm1* (Number of *acute painTRAP*^*Ai9*^ and *Oprm1* co-expressing cells/total number of TRAP ^*Ai9+*^ cells)*100%. Bars represent the mean +/− SEM for n=3 male and n=3 female TRAP2:Ai9 mice. Abbreviations: CL, central lateral. CM, central medial nucleus. IAM, interanteromedial nucleus. IMD, intermediodorsal nucleus. MDc, mediodorsal nucleus, central part. MDl, mediodorsal nucleus, lateral part. MDm, mediodorsal nucleus, medial part. PC, paracentral nucleus. PV, paraventricular nucleus. Re, nucleus reuniens. Rh, rhomboid nucleus. Xi, xiphoid nucleus. (Scale bars: 250 µM)

We also found that 75-95% of MTh neurons are nociceptive-activated **(Fig. 3G)**. 65-85% of cells express both *Oprm1* and *acute painTRAP*^*Ai9*^ **(Fig. 3H)**, with 85-100% of *Oprm1*^+^ co-expressing *acute painTRAP*^*Ai9*^ **(Fig. 3I)**, and 75-95% of *acute painTRAP*^*Ai9+*^ coexpressing *Oprm1* **(Fig. 3J)**. Notably, we did not observe differences between the left and right hemispheres in these parameters **(Fig. S4)**, reflecting previous evidence that nociceptive-activation is not lateralized in MTh. These findings, showing that most nociceptive-activated cells in the MTh express *Oprm1* and that nearly all *Oprm1*+ cells are activated by acute nociception, suggest a strong potential for opioid-mediated inhibition of nociceptive signaling at this thalamic locus.

### MTh^MOR^ cells receive widespread monosynaptic inputs

After establishing the engagement of MTh^MOR^ neurons during nociception, our next goal was to map the connectivity architecture of MTh^MOR^. First, we identified regions that send monosynaptic inputs to MTh^MOR^ by retrograde tracing with a modified rabies virus in *Oprm1*-T2A-Cre mice^50^ **(Fig. 4A)**. First, we injected a mixture of Cre-dependent TVA-mCherry and G-protein “helper” viruses to gain access to MTh^MOR^ neurons **(Fig. 4B)**. Subsequently, we transfected these cells with the GFP-tagged EnvA-pseudotyped rabies virus to visualize MTh^MOR^’s direct retrograde synaptic partners. We targeted the CL/MD nuclei specifically for our tracing experiments due to their activation during acute pain **(Fig. 2G)**, the nociceptive stability of these nuclei **(Fig. 1G)**, and their anatomical connections with nociceptive cortical neurons **(Fig. S6)**. We observed substantial monosynaptic inputs across the brain in the cerebral cortex, diencephalon, subcortical forebrain, midbrain, and nuclei of the pons, as well as the spinal cord **(Fig. S7)**. Specifically, dense inputs were found in the anterior cingulate cortex (ACC, 7.00% of total inputs), motor cortex (M2, 5.89%), habenula (Hb, 16.99%), reticular thalamus (RT, 15.22%), periaqueductal gray (PAG, 8.21%), reticular formation (RF, 7.60%), superior colliculus (SC, 12.27%), and parabrachial nucleus of the pons (PBN, 5.07%). Less dense inputs were also found from insular cortex (IC), orbitofrontal cortex (OFC), retrosplenial cortex (RSC), somatosensory cortex (S1), zona incerta (ZI), amygdala (Amy), bed nucleus of the striatal terminalis (BNST), dorsal raphe (DR), precuneiform area (PrCnF), locus coeruleus (LC), pontine reticular nucleus (PnO), pedunculotegmental nucleus (PTg), trigeminal nucleus (TGN) and the spinal cord **(Fig. 4D, I)**. Several of these regions, including the ACC^9,29^, PBN^51,52^, PAG^53,54^, S1^55^, TGN^56,57^, and spinal cord^5,11^ play crucial roles in nociceptive processing. Others like ZI^58,59^, RT^60^, and Hb^61^ are sources of inhibitory inputs to MTh^18^. Of all input regions, the ACC, PAG, PBN, RT, and RF^18^ are all either heavily implicated in nociceptive processing or regulating MTh function through modulatory or inhibitory control **(Fig. 4C)**.

**Fig. 4.**
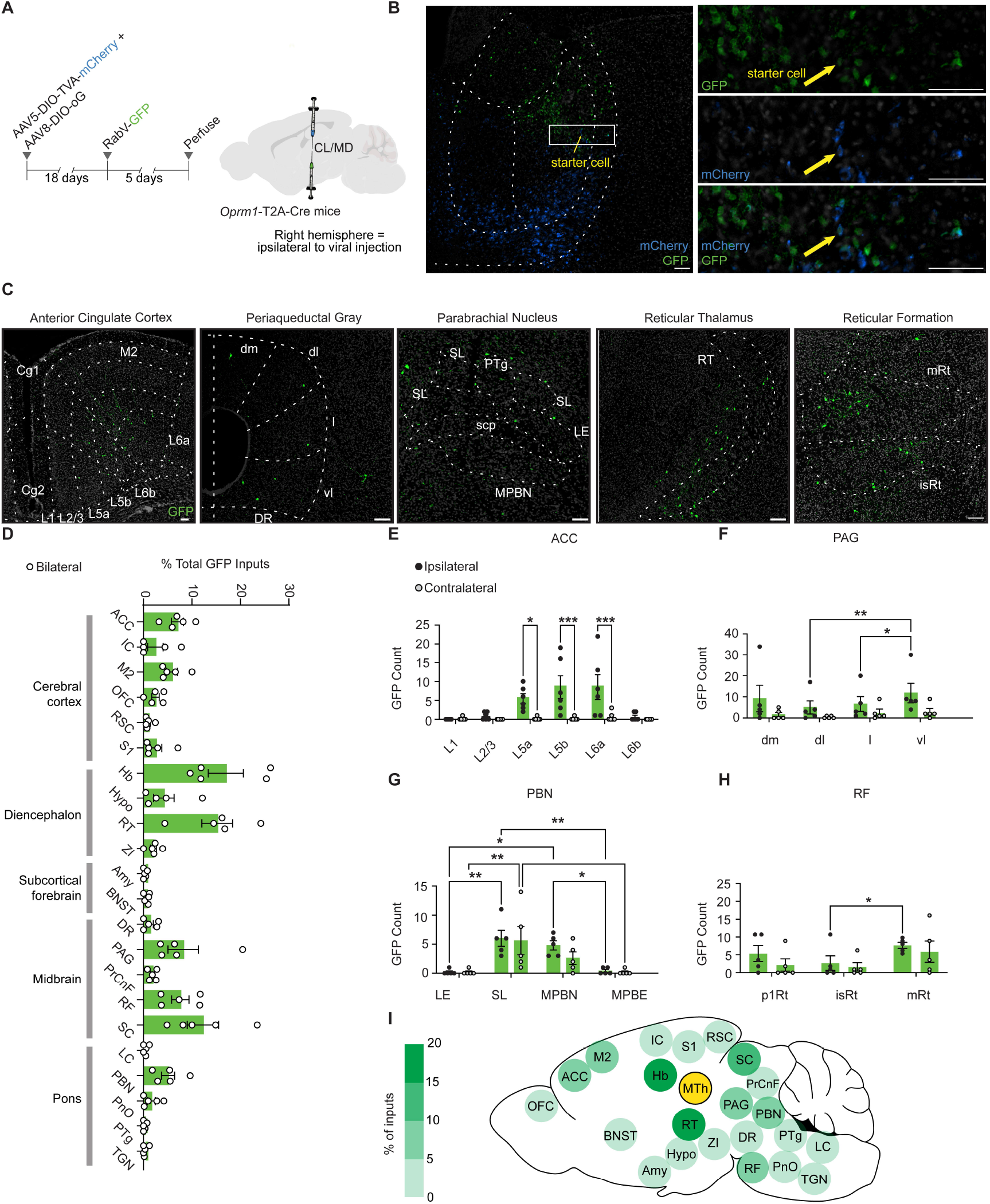
MTh^MOR^ cells receive widespread monosynaptic inputs. (A) Experimental timeline for rabies-mediated monosynaptic labeling of inputs to MOR^+^ cells in MTh. MTh^MOR^ are initially transfected with a unilateral injection of a mixture of two Cre-dependent “helper” AAVs: AAV5-DIO-TVA-mCherry (containing a modified TVA receptor) AAV8-DIO-oG (containing the rabies glycoprotein). This is followed by an injection of the EnvA-pseudotyped, G-deleted, GFP-expressing rabies virus (RabV-GFP) into the same MTh injection site in MD/CL thalamus. Ipsilateral hemisphere=same side as viral injection. (B) 20X representative image of the injection site showing expression of RabV-GFP and fluorescently labeled helper viruses in starter cells of *Oprm1*-T2A-Cre mice. (C) 20x representative images of RabV-GFP-labeled (green) cells in notable input regions, broken down by subnuclei. (B-C) (scale bars: 250 µM) (D) Quantification MTh input regions presented as percent of total inputs (Number of GFP^+^ cells in a single region/total number of GFP^+^ cells in all regions)*100% (n=3 female and n=2 male *Oprm1*-T2A-Cre mice). Bars represent the mean percentage +/− SEM; dots represent values for individual mice. (E-H) Quantification of raw number of GFP^+^ cells in hemisphere ipsilateral vs. contralateral to the injection site in subregions of ACC (n=4 female, n=2 male *Oprm1*-T2A-Cre mice), PAG (n=3 female, n=2 male *Oprm1*-T2A-Cre mice), PBN (n=3 female, n=2 male *Oprm1*-T2A-Cre mice), and RF (n=3 female, n=2 male *Oprm1*-T2A-Cre mice). Bars represent the mean percentage +/− SEM; dots represent values for individual mice. ((E-H) Two-way ANOVA with Bonferonni multiple comparisons determined significance (*p<0.05, **p<0.01, ***p<0.001)). (I) Whole-brain schematic of GFP^+^ inputs to starter cells (yellow) in MTh, visual representation of the quantification in panel (D). Abbreviations: ACC, anterior cingulate cortex (Cg1, prelimbic area. Cg2, infralimbic area). L1, cortical layer 1. L2/3, cortical layers 2/3. L5a, cortical layer 5a. L5b, cortical layer 5b. L6a, cortical layer 6a. L6b, cortical layer 6b). Amy, amygdala. BNST, bed nucleus of the stria terminalis. DR, dorsal raphe. Hb, habenula (LHb, lateral habenula. mHb, medial habenula). Hypo, hypothalamus. IC, insular cortex. LC, locus coeruleus. M2, secondary motor cortex. OFC, orbitofrontal cortex. PAG, periaqueductal gray (dm, dorsomedial column. dl, dorsolateral column. l, lateral column. vl, ventrolateral column). PBN, parabrachial nucleus (LE, external lateral nucleus. SL, superior lateral nucleus. MPBN, medial parabrachial nucleus. MPBE, external medial parabrachial nucleus). PrCnF, pre-cuneiform area. PnO, pontine reticular nucleus, oral part. PTg, pedunculopontine tegmental nucleus. RF, reticular formation (p1Rt, p1 reticular formation. mRt, mesencephalic reticular formation. isRt, isthmic reticular formation. RSC, retrosplenial cortex. S1, primary somatosensory cortex. SC, superior colliculus. TGN, trigeminal nucleus. ZI, zona incerta.

Afterwards, we sought to determine if there are any differences in inputs to MTh^MOR^ cells within subregions of the aforementioned inhibitory or nociceptive-relevant afferent regions, and found input-rich subregions in PBN, RF, and PAG. In the PBN, inputs from the ipsilateral medial PBN (MPBN) and superior lateral (SL) nucleus significantly outnumber those from the lateral external (LE) nucleus **(Fig. 4G)** by a factor of 5 on average. Similarly, contralateral SL inputs outnumber contralateral LE inputs. There are more inputs from SL than external medial parabrachial nucleus (MBPE) in both hemispheres. We also observed significantly more inputs from medial reticular thalamus than the isthmic reticular thalamus on the ipsilateral hemisphere. **(Fig. 4H)**.

Additionally, we found the PAG had a bias towards inputs from the ventrolateral column **(Fig. 4F)**, which contained significantly more Enk+ cells than the lateral and dorsolateral columns in the ipsilateral hemisphere. In contrast, no subregion-specific differences in inputs were detected across ACC layers in either Cg1 or Cg2 of both hemispheres (**Fig. S8A–I**). We next aimed to assess whether inputs to MTh^MOR^ exhibited laterality, and observed that ACC layers 5a, 5b, and 6a primarily sends ipsilateral projections to MTh-^MOR^ **(Fig. 4E)**. No hemispheric bias was observed in the other major afferent regions analyzed **(Fig. 4F; G-H)**.

### Endogenous opioid inputs are distinct from nociceptive-activated inputs to MTh

Endogenous opioids like enkephalins act on medial thalamus to mediate analgesia during stress and expectation^32–35^. Enkephalins are not likely released locally within the thalamus due to little to no enkephalinergic neurons in this site (**Fig. S10A-B**), but sources of endogenous opioid inputs that potentially act on MTh neurons have not yet been identified. To map enkephalinergic inputs to MTh, we performed retrograde tracing with a Cre-dependent viral construct in Penk-Cre mice. We injected a retrograde color switch virus that labels enkephalinergic (Enk+) inputs to MTh with GFP and non-enkephalinergic (Enk-) inputs with mCherry **(Fig. 5B)** and identified enkephalinergic inputs across the brain, with a large density in the habenula, anterior cingulate cortex, periaqueductal gray, retrosplenial cortex, and parabrachial nucleus of the pons **(Fig. 5A)**.

**Fig. 5.**
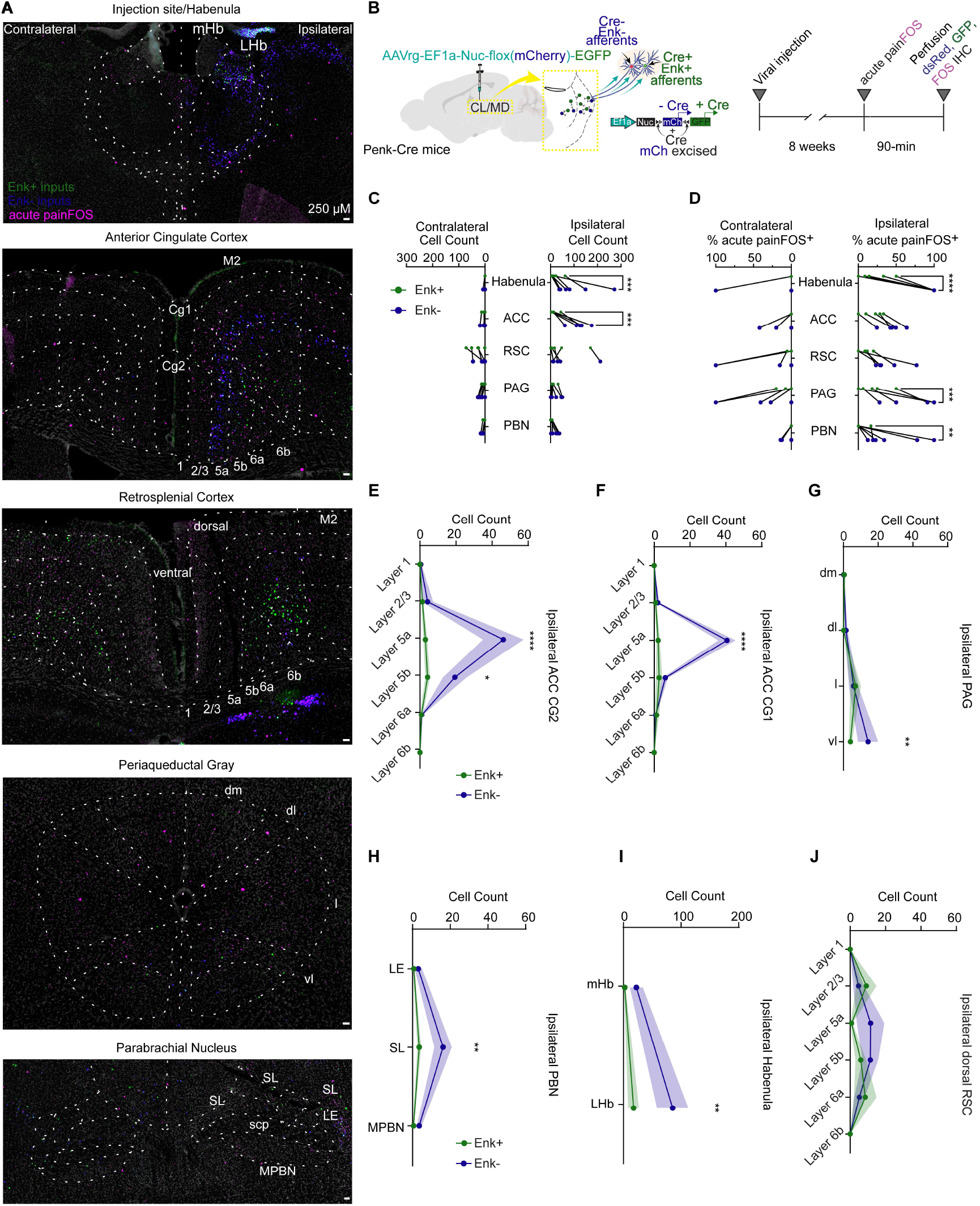
Endogenous opioid inputs are distinct from nociceptive inputs to MTh. (A) 20x representative images of color switch virus-labeled inputs to MTh. Top image shows injection site and habenula inputs. Subsequent images show bilateral inputs to MTh from ACC, RSC, PAG, and PBN. Green= enkephalinergic inputs to MTh (Enk+). Blue= non-enkephalinergic inputs to MTh (Enk-). Magenta= 55C hot water stimulus-activated cells (acute painFOS). (scale bars: 250 µM). (B) Experimental timeline for expression of AAVrg-EF1α-Nuc-flox(mCherry)-EGFP (AKA color switch virus) in CL thalamus of Penk-Cre mice followed by acute painFOS protocol. In the presence of Cre, mCherry will be deleted, labeling Cre+ (AKA enkephalinergic) cells with GFP. In the absence of Cre, mCherry is not deleted, labeling Cre- (AKA non-enkephalinergic) cells with mCherry. (C) Quantification of number of Enk+ vs Enk-inputs to MTh. (D) Quantification of the percent of acute painFOS^+^ neurons co-expressing GFP (green) or mCherry (blue) (Number of GFP-expressing cells co-expressing acute painFOS)/total number of GFP-expressing cells)*100%. Number of mCherry-expressing cells co-expressing FOS)/total number of mCherry-expressing cells)*100%. (E-J) Quantification of raw number of GFP-expressing and mCherry-expressing cells in subregions of ACC, PAG, PBN, Habenula, and RSC. Points represent mean +/− SEM. ((C-J) Two-way ANOVA with Bonferonni multiple comparisons determined significance (*p<0.05, **p<0.01, ***p<0.001 ****p<0.0001)) on n=6 male *Penk*-Cre mice). Ipsilateral=hemisphere on the same side as acute painFOS stimulus. Contralateral=hemisphere on the opposite side of acute painFOS stimulus). Abbreviations: ACC, anterior cingulate cortex (Cg1, prelimbic area. Cg2, infralimbic area). L1, cortical layer 1. L2/3, cortical layers 2/3. L5a, cortical layer 5a. L5b, cortical layer 5b. L6a, cortical layer 6a. L6b, cortical layer 6b). CL, central lateral nucleus of the thalamus. Habenula (LHb, lateral habenula. mHb, medial habenula). PAG, periaqueductal gray (dm, dorsomedial column. dl, dorsolateral column. l, lateral column. vl, ventrolateral column). PBN, parabrachial nucleus (LE, external lateral nucleus. SL, superior lateral nucleus. MPBN, medial parabrachial nucleus).

First, we investigated whether Enk+ inputs to MTh outnumbered Enksources and observed that in the habenula and ACC on the hemisphere ipsilateral to the viral injection, there were significantly more Enkinputs than Enk+ inputs **(Fig. 5C)**. Subregion-specific analyses revealed significantly more Enk-than Enk+ neurons in Layers 5a and 5b of the ipsilateral ACC Cg2, and in layer 5a of ipsilateral Cg1 of ACC **(Fig. 5E-F)**. Additionally, in the hemisphere ipsilateral to the injection, the ventrolateral PAG, superior lateral PBN, lateral habenula contained significantly more Enkthan Enk+ cells **(Fig. 5G-I)**, while other subnuclei of these areas did not show significant differences between these cell types **(Fig. 5J; S9A-D, F, H)**. In the contralateral hemisphere, the ventrolateral PAG had significantly more Enk-than Enk+ cells by a factor of 3.5 on average **(Fig. S9G)**. Notably, Layer 5b of the contralateral RSC had significantly more Enk+ cells than Enk-neurons. RSC was the only quantified brain area in which Enk+ neurons outnumbered their non-enkephalinergic counterparts **(Fig. S9E)**.

Along with their expression of enkephalin, the ACC^9,29^, PAG^53,54^, RSC^62^, and PBN^51,52^ also critically mediate nociceptive processing. It is not known whether these enkephalinergic inputs also project nociceptive information to MTh. To determine whether the nociceptive-activated neurons in these input areas are the same ones expressing enkephalin, we quantified overlap of acute painFOS and Enk+ or Enk- and found that in the ipsilateral habenula, PAG, and PBN, the Enk-inputs were significantly more likely to be noci-ceptive-activated **(Fig. 5D)**. Similar results were found in the ACC and RSC, but were trending and not significant. These results indicate that nociceptive-active circuits in these brain areas are distinct from enkephalinergic inputs to MTh.

### MTh^MOR^ neurons send widespread outputs throughout the brain

Previous literature shows the connections with MTh nuclei are often reciprocal, mediating bidirectional communication between MTh and other brain regions^18,63,64^. To identify which of our input regions engage MTh^MOR^ reciprocally, we mapped the brain-wide outputs from MTh^MOR^. We leveraged viral-mediated circuit tracing to map MTh^MOR^ outputs by using a Cre-dependent mCherry-tagged anterograde virus into MTh of *Oprm1*-T2A-Cre mice **(Fig. 6A-B)**.

**Fig. 6.**
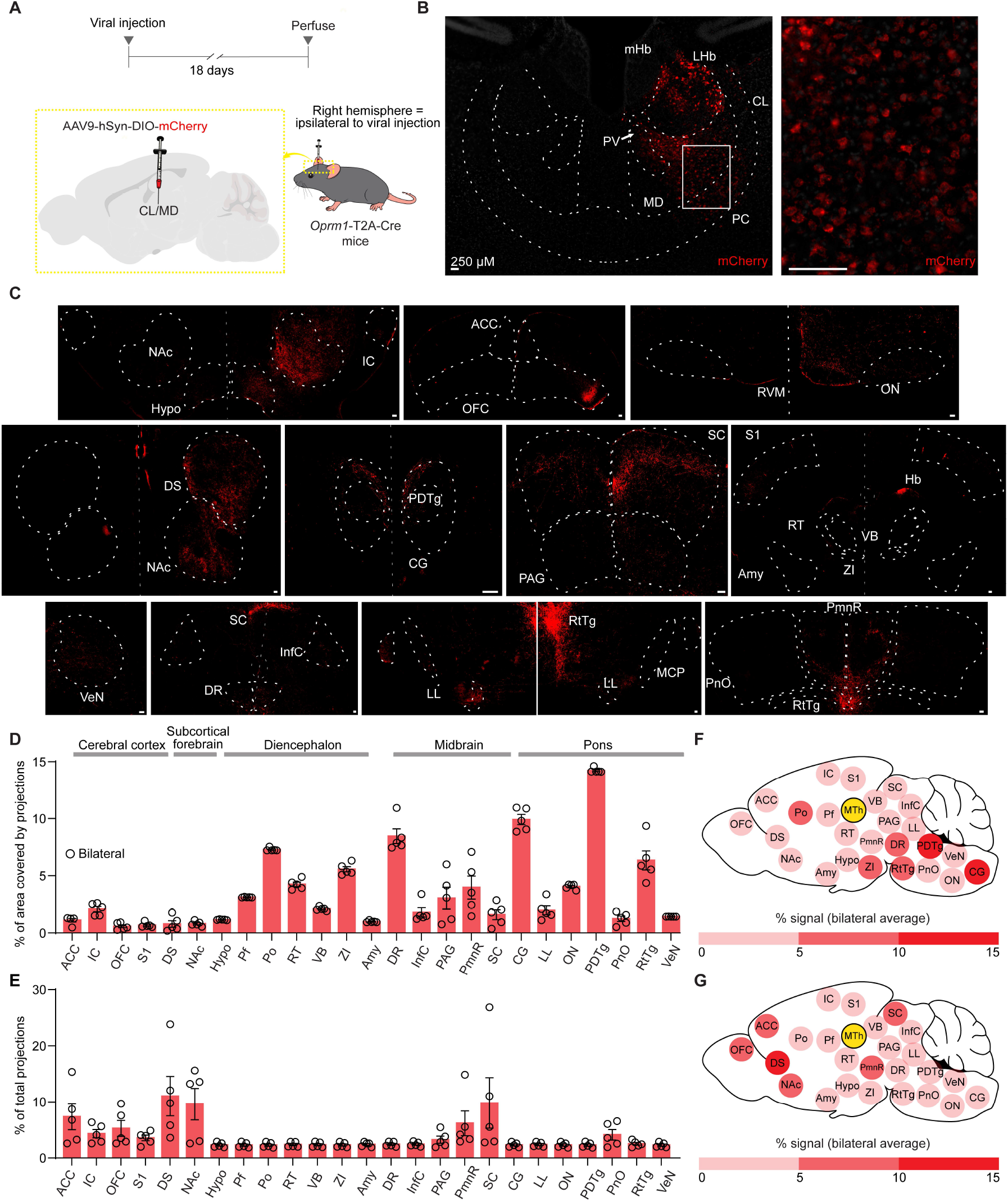
MTh^MOR^ cells send widespread outputs. (A) Experimental timeline labeling of outputs from MOR^+^ cells in MTh. MTh is transfected (with needle targeting CL/MD thalamus) with a Cre-dependent, mCherry-tagged fluorescent virus in *Oprm1*-T2A-Cre mice. (B) 20x representative image of the injection site showing expression of AAV9-hSyn-DIO-hm4D(Gi)-mCherry virus in MTh. (C) 20x representative images of MTh^MOR^ axons in notable output regions, broken down by subnuclei. Red=mCherry. (B-C) scale bars: 250 µM). (D) Quantification of MTh^MOR^ outputs presented as percent of each area covered by MTh^MOR^ projections (n=3 female and n=3 male *Oprm1*-T2A-Cre mice). (E) Quantification of MTh^MOR^ outputs presented as percent of total projections (Number of pixels per region/total number of pixels across regions)*100%. (F) Whole-brain schematic of MTh^MOR^ outputs, visual representation of the quantification in panel (D). (G) Whole-brain schematic of MTh^MOR^ outputs, visual representation of the quantification in panel. (E). (D-G) Bars represent the average across left and right hemispheres for each region; dots represent values for individual mice. Abbreviations: ACC, anterior cingulate cortex. Amy, amygdala. CG, central gray. DR, dorsal raphe. DS, dorsal striatum. Hypo, hypothalamus. IC, insular cortex. InfC, inferior colliculus. LL, lateral lemniscus. NAc, nucleus accumbens. OFC, orbitofrontal cortex. ON, olivary nucleus. PAG, periaqueductal gray. PDTg, posterodorsal tegmental nucleus. Pf, parafascicular nucleus of the thalamus. PmnR, paramedian raphe. PnO, pontine reticular nucleus, oral part. RT, reticular nucleus of the thalamus. RtTg, reticulotegmental nucleus. S1, primary somatosensory cortex. SC, superior colliculus. VB, ventrobasal complex of the thalamus. VeN, vestibular nucleus. ZI, zona incerta.

We observed projections throughout the cerebral cortex, diencephalon, subcortical forebrain, midbrain, and pons. MTh^MOR^ projects to the anterior cingulate cortex (ACC), amygdala (Amy), central gray (CG), dorsal raphe, dorsal striatum (DS), hypothalamus, insular cortex, inferior colliculus (InfC), lateral lemniscus (LL), nucleus accumbens (NAc), orbitofrontal cortex, olivary nucleus (ON), periaqueductal gray, posterodorsal tegmental nucleus (PDTg), parafascicular nucleus of the thalamus (Pf), paramedian raphe (PmnR), pontine reticular nucleus, reticular nucleus of the thalamus, reticulotegmental nucleus (RtTg), primary somatosensory cortex, superior colliculus, ventrobasal complex of the thalamus (VB), vestibular nucleus (VeN), and the zona incerta **(Fig. 6C)**.

Using the AxoDen automated platform to quantify axon density^65^, we found that 1-15% of these areas were covered by MTh^MOR^ projections **(Fig. 6D,F)**, and most areas received a very similar amount of projections, with some regions standing out with a larger proportion of axonal projections: ACC (7.34%), IC (4.32%), OFC (5.26%), DS (11.03%), NAc (9.60%), PmnR (6.18%), and SC (9.82%) **(Fig. 6E,G)**. Together, these extensive connections of MTh^MOR^ neurons, many of which are bidirectional in nature, highlight the importance of this population in brain-wide communication, and its connections with areas also involved in nociception further implicate its role in nociceptive processing.

## Discussion

Here, we identify a highly connected subpopulation of MOR+ neurons in the medial thalamus (MTh^MOR^) that are robustly activated by nociceptive stimuli and may remain stably engaged across both acute and chronic pain states. This work establishes a foundation for understanding how opioidergic signaling modulates thalamic nociceptive circuits and highlights MTh^MOR^ neurons as a promising target for therapeutic intervention in pain. Beyond their role in nociception, mapping these neurons contributes to the creation of a new thalamic atlas of nociceptive and opioidergic cell types, providing a framework for exploring the broader functions of the MTh in sensory, affective, and cognitive processes. Future studies delineating the role of MTh^MOR^ neurons in non-nociceptive behaviors will be essential for guiding the development of selective and effective analgesic strategies.

### Non-uniform distribution of nociceptive responses in MTh

The present study is the first ever to assess nociceptive responses in all ten MTh subnuclei at once, and supports the previous but limited evidence that MTh does not act a as a singular functional unit. Past studies suggest MTh is a functionally heterogenous structure^19,64^ whose subnuclei respond differently to noxious stimuli such as cold exposure, pinch, and electrical stimulation of the sciatic nerve. For instance, noxious cold evokes elevated FOS expression in Xi^23^, unlike other MTh nuclei, and sciatic nerve stimulation produces different amplitudes of evoked activity between CL, PC, and MD thalamus^24^. Noxious pinch even reduces activity of some MTh units^66^, while activating others.

Our whole-nucleus analysis reveal elevated responses to acute nociceptive stimuli in CM, IAM, IMD, Re, and Xi, which are notably all located along the midline. The midline thalamus has long-since been implicated in processing nociceptive stimuli^4,10,13,67,68^, and display strikingly different afferent/efferent connections^64^ and genetic profiles^69^ from other MTh subnuclei^19^, which could explain this biased activation.

However, our slice-by-slice investigation further reveals discrete acute nociceptive hotspots within the spatial axes of multiple MTh nuclei including CL, MD, PC, PV, and Rh. These observations are consistent with prior work demonstrating localized nociceptive processing within these regions. In particular, we identify a nociceptive hotspot within the anterior PV, paralleling previous findings that show anterior PV is specifically recruited during somatic^70^ and visceral^71^ pain hypersensitivity.

### A MTh ensemble that is stable across pain states

Our activity-dependent labeling of nociceptive responses in MTh reveals the presence of a subpopulation across multiple MTh nuclei that was engaged during multiple acute pain experiences and during allodynia in chronic pain. Previous work from our lab has identified similar ensembles that are stable across pain states in the BLA^47,48^ and ACC^29^. What does it mean for a thalamic ensemble to be stably activated across acute pain states? First we must consider that the activation of any thalamic neuron during nociception could be coincidental, due to the fact that the thalamus is tonically active during baseline, wakeful states^72–74^. However, if that neuron’s activity is elevated compared to home cage conditions, and especially if it is activated during two of the same acute nociceptive stimuli given at temporally distinct points, repeated engagement could be evidence towards this cell being critical for nociceptive processing in the area.

On the other hand, repeated engagement could also reflect learning/memory processing. The initial activation could represent the surprise of a highly salient noxious stimulus, while the subsequent activation could be the re-activation of a pain memory, or a representation of a conditioned stimulus response. Conditioned responses to noxious stimuli are encoded within the thalamus^19^, specifically in MD^75^ and Re^76^. Another explanation to a neuron being responsive to pain at multiple time points could be that it is a reliably pain-projecting relay neuron. Due to the thalamus’ connectivity with the ACC^9^, as well as its established necessity for transmitting affective nociceptive information to this region^24^, these pain-stable MTh neurons could be critical to relaying affective nociceptive information to the cortex.

What would it mean for an MTh ensemble to be stably activated during acute and chronic pain? It would be easy to assume that this ensemble is critical for processing and relaying both acute and nociceptive information. However, the medial thalamus is also heavily involved in other functions such as arousal^16,19,74,77–79^, as well as directing attention to particular stimuli^18,19,77,80,81^. Therefore, it could be that MTh neurons that are activated during evoked acute and chronic pain stimuli due to the highly salient nature of these stimuli. They could be recruited to help draw attention to these incoming noxious, distracting stimuli, rather than to simply relay affective nociceptive information. Others have also posited that given the presence of a separate system of lateral thalamocortical relay circuits for projecting specific nociceptive information, it is likely that medial thalamus facilitates attentional engagement and affective qualities of noxious stimuli rather than directly contributing to sensory perception, for acute and chronic pain^18,82^.

### MTh activity increases during chronic pain-induced mechanical allodynia

Murine and human studies have also observed elevated activity in MTh nuclei during chronic pain states. The mouse PV exhibits increased FOS expression during SNI and Complete Freud’s Adjuvant (CFA)-induced persistent pain states. MD exhibits increased spontaneous firing after a spinal cord injury (SCI) in rats^83^, which is postulated to be a result of reduced inhibitory control from the zona incerta. These investigators also found that a noxious pinch in the hypersensitive hind paws further elevates MD activity in SCI and not sham mice^83^. In another study, PV displays increased FOS expression after colorectal distension in a chronic visceral pain model^84^. Medial thalamic nuclei are also hyperactive in humans with deafferentation pain^85^.

### Role of MOR in nociceptive MTh and potential role of enkephalin

It has long-since been established that MORs are densely expressed in MTh^37,50^, and that MTh^MOR^ plays a crucial role in the affective component of nociception^20^. Other brain regions also exhibit high overlap of nociceptive neurons with MOR^+^ cells^2,29^. Additionally, it has been shown *ex vivo* and *in vivo* that opioid agonists reduce MTh activity^24,39,86^, indicating that MOR agonists inhibit MTh activity and in turn inhibit nociceptive behavioral responses. Therefore, MOR-targeted analgesics could be developed to target MTh for pain.

However, exogenous opioid administration can lead to a host of harmful side effects such as respiratory depression, addiction, and rebound hyperalgesia^40–42^. On the other hand, endogenous opioids are released in small amounts at specific sites throughout the nervous system to induce analgesia in a controlled manner during placebo analgesia and stress-induced analgesia in a way that avoids these side effects^2^. Notably, these two types of endogenous pain relief have been shown to specifically reduce MTh activity^32,35,49^, but it remains unknown whether this reduction is due to release of endogenous opioids directly onto MTh^MOR^ neurons, or an artifact of the blockade of ascending nociceptive information from the spinal cord, which is traditionally known as the primary locus of the endogenous analgesic response^87^.

Interestingly, there is evidence that the endogenous opioid enkephalin could act directly on MTh^MOR^ neurons. Enkephalinergic axons from the PBN^88^ and spinal cord^89^ have been shown in the MTh. Additionally, the vlPAG projects to MTh^13^ and also expresses enkephalin^90^, though the overlap between these populations has not been proven until the present study, which quantifies enkephalinergic inputs from vlPAG to MTh. The present study also identifies ACC, RSC, and habenula as enkephalin-expressing inputs to MTh; these regions are afferents to MTh^18,61,91^, involved in nociceptive processing^9,24,61,62^, and the ACC in particular has been shown in murine and human studies to play a critical role in endogenous opioid analgesia^27,33,34^. Further studies proving the direct inhibition of MTh^MOR^ by enkephalin during endogenous analgesic responses, or demonstrating that the release of enkephalin onto MTh mediates analgesia *in vivo*, may reveal a potential enkephalin-mediated circuit-based analgesic therapy for chronic pain. Notably, enkephalin is primarily an agonist for the delta opioid receptor^92,93^, so further characterization of DOR in nociceptive MTh neurons will also be necessary.

### Brain-wide connectivity with MTh^MOR^

The highly interconnected nature of MTh^MOR^ with brain areas critical for nociception^10,94,95^ supports its originally known function as an affective relay center for projecting non-specific nociceptive information diffusely across the cortex^96^. However, its connectivity with regions outside of the pain matrix reinforces its importance in non-nociceptive functions as well like action selection^97^, arousal^79^, attention^19^, and cognition^18,80^.

We next compared the connectivity architecture of MTh^MOR^ with the previously-known connectivity of the broader population of MTh neurons. The present investigation is consistent with previous anatomical evidence that the MTh receives widespread inputs and outputs^18,19^. However, the MOR^+^ population displays unique inputs from habenula and precuneiform area and outputs to inferior colliculus, central gray, lateral lemniscus, vestibular nucleus, and olivary nucleus. We also did not observe parietal cortex, perirhinal cortex, piriform cortex, parasubthalamic nucleus, suprachiasmatic nucleus, SNr, subiculum, tuberomammilary nucleus, visual cortex afferents/efferents that have been reported in the literature^80^. It is possible that the connectivity of the opioidergic MTh neurons displays features distinct from that of the broader region. It is also possible that our tracers, which bled into MOR^+^ habenular neurons adjacent to the MTh, could explain this, especially as connectivity with the habenula has been established in several of these areas, including the SCN and SNr^98,99^.

## Limitations and future directions

It is important to understand that the presence of nociceptive-activated cells does not imply the concomitant existence of nociceptive-specific cells in the MTh, which would be activated uniquely by nociceptive stimuli. The medial thalamus does not encode specific information about sensory stimuli-that is reserved for lateral thalamic nuclei^9,96^. Due to this feature, it is unlikely that nociceptive-specific cells exist in MTh. However, investigating nociceptive-activated neurons in MTh can still provide valuable information about how MTh processes noxious stimuli and brings these stimuli into our attention for proper action selection.

Our study also relies completely on FOS-based tools to label active neurons. This is a static measure lacking the real-time activity dynamics that could be captured with *in vivo* imaging of these cells during pain. Additionally, FOS is not a perfect marker of neural activityit is produced in non-neuronal cellsand is not made by all activated neurons^100^. Future studies that utilize techniques like *in vivo* electrophysiology or calcium imaging are better suited to tease out the temporal nuances of the differential responses of MTh neurons to nociceptive states.

Additionally, further work is needed to assess the behavioral impact of selective inhibition of MTh^MOR^ during pain to further prove the necessity of this cell-type for nociceptive responses. A previous study that microinjected a MOR agonist during pain^20^ indicates that MTh^MOR^ are necessary for pain processing, but lacks chronic pain modalities, as well as a comprehensive read-out of pain responses, only relying a couple measures of behavioral responses to pain.

In conclusion, this study establishes a comprehensive framework for understanding how nociceptive and opioidergic signaling converge in the medial thalamus. By integrating large-scale anatomical, molecular, and circuit analyses, we reveal a stable and highly connected MTh^MOR^ population that serves as a nexus between nociceptive input and affective-motivational processing. These findings provide a foundation for a new thalamic atlas of nociceptive and opioidergic cell types, positioning the MTh as both a key relay in endogenous analgesic circuits and a promising target for next-generation, circuit-specific pain therapies. Future work combining in vivo imaging, activity manipulation, and behavioral analysis will be critical to determine how MTh^MOR^ neurons shape perception, motivation, and attention in both pain and non-pain contexts—advancing our capacity to harness endogenous opioid systems for safe, precise, and durable relief from chronic pain.

## Acknowledgements

This work was supported by: National Institute of Health (NIH) awards NIGMS DP2GM140923 (G.C.), NINDS F99NS135765 (L.L.E.), NIGMS DP2GM140923-01S1 (L.L.E., G.C.), NIDA R01DA054374 (G.C.), NINDS R01NS130044 (G.C.), NIDA F31DA057795 (L.M.W.), NINDS F31NS125927 (J.A.W.), NIDA T32DA028874 (G.S.). We thank Dr. Julie Blendy for the *Oprm1*-T2A-Cre mice, the Penn University Laboratory Animal Resour-sources (ULAR) veterinarians and husbandry staff, and Dr. Kevin Beier for the custom rabies virus.

## Author contributions

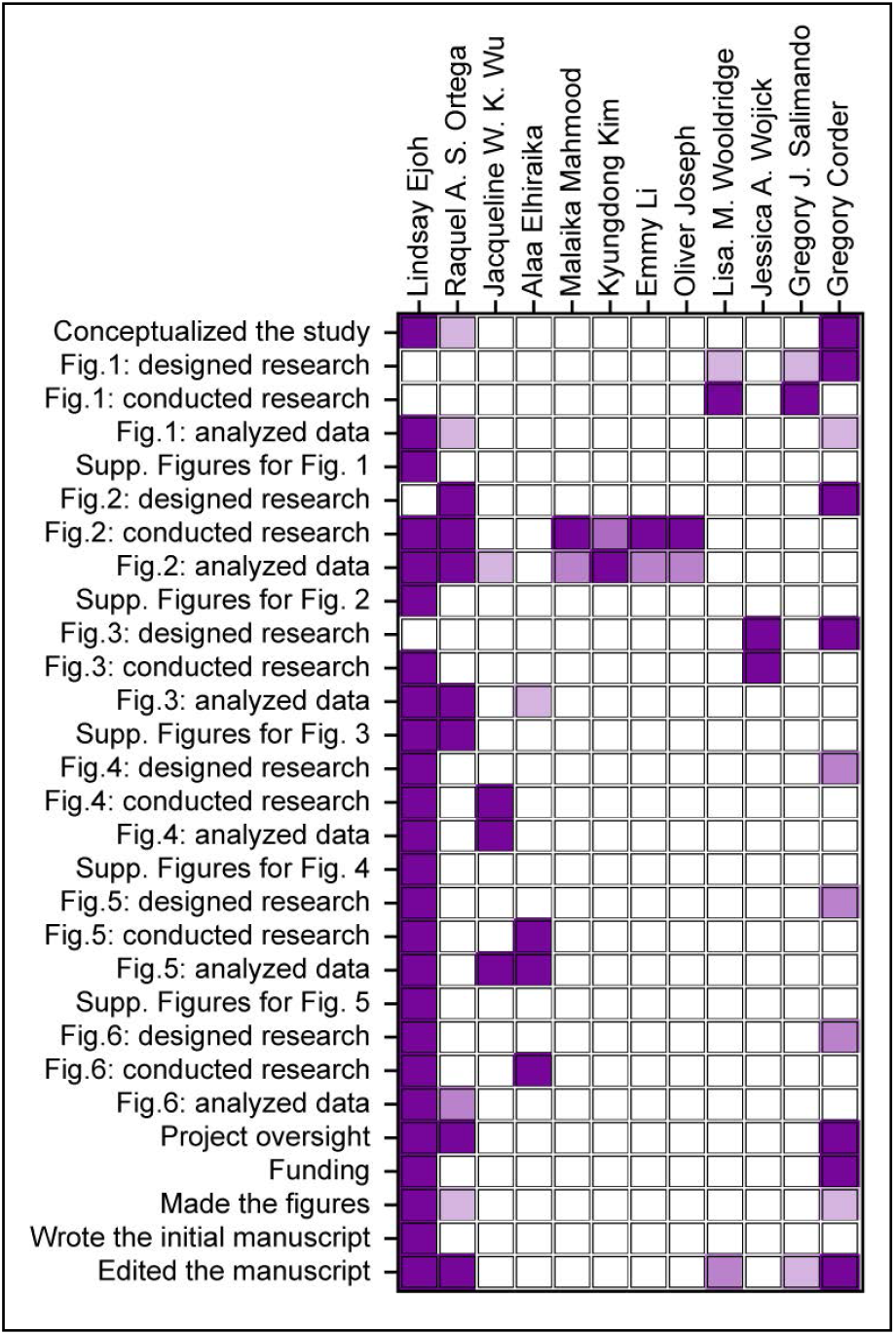

## Supplementary Materials

## Methods

### Experimental Model Details

All experimental procedures were approved by the University of Pennsylvania Institutional Animal Care and Use Committee (IACUC) to conform to US National Institutes of Health (NIH) guidelines. Both male and female transgenic mice >P50 were used in this study. All animals were drug and test-naïve, housed 1-5 per cage, under consistent temperature, humidity, and light conditions (12-hour reverse light-dark cycle, lights on from 21:00 to 09:00) and given food and water ad libitum. All experiments were performed during the dark cycle. For anatomical experiments, we utilized C57BL6/J mice, *FOS-*FOS-2A-iCreERT2 or “TRAP2” mice (FOStm2.1(iCre/ERT2)Luo) Luo/Jackson Laboratory, stock #030323) crossed with B6.Cg-Gt(ROSA)26Sor^t,9(CAG-tdTomato)Hze^/J (Ai9), TRAP2 mice crossed with a Sun1 reporter that expressed a GFP fluorophore in a Cre-dependent manner in the nuclear membrane (“TRAP2;Sun1-sfGFP”) bred to homozygosity for both genes, (Jax, stock #007914) mice bred to homozygosity in both alleles, B6;129S-Penk^tm2)(cre)Hze^/J (Penk-IRES2-Cre) mice bred to heterozygosity, and C57/BL/6NTAC-^Oprm1em1(cre)Jabl)^Mmnc (*Oprm1*-T2A-Cre) bred to heterozygosity. *Oprm1*-T2A-Cre was generated from Dr. Julie Blendy’s laboratory and bred in-house.

### Quantification and Statistical Analysis

All data are reported as mean values +/− the standard error of the mean (SEM) unless otherwise specified. Statistical analysis was performed in GraphPad Prism (USA) (Supplementary Table 1). For all statistical tests, *p<0.05, **p<0.01, ***p<0.001, and ****p<0.0001. All data was tested for normality using the Kolgomorov-Smirnoff test. The means of different data distributions were compared using One-Way ANOVA, 2-way ANOVA, Bonferonni’s multiple comparisons, mixed effects model with Bonferonni multiple comparisons, and Tukey’s multiple comparisons. Statistics details for supplemental figures are also detailed in supplemental figure legends.

### Antibodies

Chicken anti-GFP abcam ab13970 Lot GE3361051-17 1:2000 Rabbit anti-dsRed Takara 632496 Lot 2210019 1:2000 Guinea pig anti-FOS Synaptic systems 226308, Lot 1-10 1:1000 Rabbit anti-MOR abcam ab234054, Lot 1000560-11 1:500 Rabbit anti-Met-enkephalin Millipore AB1975 Lot 3821706 1:1000 Donkey anti-rabbit Alexa 594 Invitrogen Thermo Fisher A21207 Lot 2210019 1:500

Donkey anti-rabbie Alexa 555 Invitrogen A31572 Lot 2492963 1:500

Goat anti-chicken Alexa 488 Invitrogen Thermo Fisher A11039, Lot 2079383 1:500

Donkey anti guinea pig Alexa 647 Jackson Immuno Research 706-605-148 1:500

### Viruses

*AAV5-CAG-FLEX-TCB*. Titer: 1.6E13 vg/mL. Volume: 150nL. MD coordinate: AP: −1.6mm; ML: −.65mm; DV= −3.6mm

*AAV8-CAG-FLEX-RabV-G*. Titer: 2.4E12 vg/mL. Volume: 150nL. MD coordinate: AP: −1.6mm; ML: −.65mm; DV= −3.6mm

*RabV-GFP*. Titer: 6.2E8 vg/mL. Volume: 300nL. MD coordinate: AP: −1.6mm; ML: −.65mm; DV= −3.6mm from Bregma

*AAVrg-EF1a-Nuc-flox(mCherry)-EGFP*. Titer: 2.5E13 vg/mL Lot: v81646 Volume: 300nL. CL coordinate: AP: −1.34mm; ML: −0.75mm; DV= −3.4mm from Bregma

*AAV9-hSyn-DIO-hm4D(Gi)-mCherry*. Titer: 2.30E+12 vg/mL Lot: v66706 Volume: 300nL. MDl Coordinate: AP: −1.6mm; ML: −.65mm; DV= −3.6mm from Bregma

*AAV9-hSyn-DIO-mCherry*. Titer: 2.10E+13 vg/mL Lot: v75884 Volume: 500nL. ACC Coordinate: AP: +1.2mm; ML: +0.3mm; DV= −1.8mm from Bregma

### Chemicals

4-hydroxytamoxifen (4-OHT; HelloBio, HB6040)

### Mouse Models

C57BL6/J (Jackson Labs, Stock 664)

*Oprm1*-T2A-Cre (Dr. Julie Blendy, Penn)

*Penk*-IRES2-Cre mice (Stock 025112)

*TRAP2*(+/+), Fos^tm2.1(icre/ERT2)Luo)^ (Jackson Labs, Stock 030323)

*TRAP2*:Ai9(+/+, +/+), Fos^tm2.1(icre/ERT2)Luo)^ crossed with Ai9 (B6.Cg-*Gt(ROSA)26Sor*^*tm9(CAG-tdTomato)Hze*^ (Jackson Labs, Stock 7909)

*TRAP2*;Sun1-sfGFP(+/+,+/+), (FOS^tm2.1(icre/ERT2)Luo^ crossed with CAG-Sun1/sfGFP (B6.129-Gt(ROSA)26Sor^tm5.1(CAG-Sun1/sfGFP)Nat/Mmbe^ (Jackson Labs Stock 30952)

### Softwares and Algorithms

HALO-AI for automated cell quantification

AxoDen for automated axon quantification

Adobe Illustrator for anatomical border drawings

GraphPad Prism for statistical analyses and data visualization

### Immunohistochemistry (Brain tissue)

Mice were anesthetized with FatalPlus (Vortech) and then transcardially perfused with 0.1 M phosphate buffered saline (PBS) and 10% normal buffered formalin (NFG, Sigma, HT501128). Brains were removed and post-fixed in 10% NBF for 18 hours at 4°C, then placed in a 30% sucrose in PBS solution for 48 hours until they sunk to the bottom of their storage tubes. Afterwards, brains were frozen in O.C.T. (Thermo Fisher Scientific) at −20C before being sectioned coronally at 30 µM and placed free-floating in PBS. When required, a small incision was made to cortex to denote side contralateral to viral injection. Immunostaining of tissue occurred within 2-14 days of sectioning and every step occurred at room temperature. First, sections were permeabilized in 0.3% Triton X-100 solution in PBS (PBS-T). Then, the tissue was blocked in a solution of 5% Normal Donkey Serum (NDS) in 0.3% PBS-T for 2 hours to prevent non-specific binding of antibodies before soaking in a solution of the primary antibody diluted in blocking solution for ∼18 hours. After 3 additional PBS-T washes, brain sections were soaked in secondary antibody diluted in the blocking solution for 2 hours. After 3 additional PBS-T washes and 2 PBS washes, nuclei were stained with DAPI solution diluted 1:10,000, mounted on positively-charged Superfrost Plus glass slides (Fisher Scientific, 1255015) and cover slipped with 24×50mm glass coverslips (WVR, L-WVR-CG-R4) and Fluromount-G mounting medium (Invitrogen, 00-4958-02).

### Immunohistochemistry (Spinal cord tissue)

Mice were anesthetized with FatalPlus (Vortech) and then transcardially perfused with 0.1 M phosphate buffered saline (PBS) and 10% normal buffered formalin (NFG, Sigma, HT501128). Spinal cords were dissected and post-fixed in 10% NBF for ∼2 hours at RT, then placed in a 30% sucrose in PBS solution until they sunk to the bottom of their storage tubes. Afterwards, spinal cords were frozen in O.C.T. (Thermo Fisher Scientific) at −20C before being sectioned transversely at 30 uM and placed free-floating in PBS.

Immunostaining of tissue occurred within 14-21 days of sectioning and every step occurred at room temperature. First, sections were permeabilized in 0.3% Triton X-100 solution in PBS (PBS-T). Then, the tissue was blocked in a solution of 5% Normal Donkey Serum (NDS) in 0.3% PBS-T for 2 hours to prevent non-specific binding of antibodies before soaking in a solution of the primary antibody diluted in blocking solution for ∼18 hours. After 3 additional PBS-T washes, spinal cord sections were soaked in secondary antibody diluted in the blocking solution for 2 hours. After 3 additional PBS-T washes and 2 PBS washes, nuclei were stained with DAPI solution diluted 1:10,000, mounted on positively-charged Superfrost Plus glass slides (Fisher Scientific, 1255015) and cover slipped with 24×30mm glass coverslips (WVR, 48404-467) and Fluromount-G mounting medium (Invitrogen, 00-4958-02).

### FOS induction

#### acute painFOS

We habituated mice to a testing room for two to three consecutive days. During these habituation days, no nociceptive stimuli were delivered and no baseline thresholds were measured (i.e. mice were naïve to pain experience before the FOS procedure). We placed individual mice within red plastic cylinders (∼9-cm D), with a red lid, on a raised perforated, flat metal platform (61-cm x cm-26). The experimenter’s sat in the testing room for the thirty minutes of habituation; this was done to mitigate potential alterations to the animal’s stress and endogenous antinociception levels to the presence of experimenter, behavior room, and Von Frey rack. To execute FOS induction, we placed mice in their habituated cylinder for 30 min, and then a 55°C water droplet (water was heated to 70C and rapidly cooled upon application) was applied to the central-lateral plantar pad of the left hindpaw) once every 30-sec for 10 min. Following the hot water stimulations, the mice remained in the cylinder for an additional 90 min before being injected with FatalPlus intraperitoneally and transcardially perfused.

#### controlFOS

We habituated each mouse to its own home-away-from-homecage for 2 consecutive days. On day 3, at least 2 hours into the dark cycle, mice were gently removed from their home cages and placed in home-away-from-home-cage for 90 minutes before administering FatalPlus injections and perfusions.

### TRAP protocol

#### acute painTRAP

Acute painTRAP induction was performed similarly to previous studies^47^. We habituated mice to a testing room for two to three consecutive days. During these habituation days, no nociceptive stimuli were delivered and no baseline thresholds were measured (i.e. mice were naïve to pain experience before the TRAP procedure). We placed individual mice within red plastic cylinders (∼9-cm D), with a red lid, on a raised perforated, flat metal platform (61-cm x 26-cm). The experimenter’s sat in the testing room for the thirty minutes of habituation; this was done to mitigate potential alterations to the animal’s stress and endogenous antinociception levels to the presence of experimenter, behavior room, and Von Frey rack. To execute the TRAP procedure, we placed mice in their habituated cylinder for 30 min, and then a 55°C water droplet (water was heated to 70C and rapidly cooled upon application) was applied to the central-lateral plantar pad of the left hindpaw) once every 30 s for 10 min. Following the water stimulations, the mice remained in the cylinder for an additional 60 min before injection of 4 hydroxytamoxifen (4-OHT, 40 mg/kg in vehicle; subcutaneous). After the injection, the mice remained in the cylinder for an additional 4 hours to match the temporal profile for FOS expression, at which time the mice were returned to the home cage.

#### controlTRAP

We habituated each mouse to its own home-away-from-homecage for 2 consecutive days. On the last day, at least 2 hours into the dark cycle, mice were gently removed from their home cages and placed in home-away-from-home-cage. Mice were then injected with 4-OHT (40 mg/kg in vehicle; subcutaneous) and returned to their home cages.

### Stereotaxic surgical injections

Adult mice (at least 8 weeks of age) were anesthetized with isoflurane gas in oxygen (initial dose 5%, maintenance dose 1.5-2%), and fitted into a Kopf stereotaxic frame for all surgical procedures. After checking for proper surgical depth and disinfecting the head with 70% ethanol and betadine, an incision was made in the skin to expose the skull. Then, a hole was drilled before 10 uL Nanofil Hamilton syringes (WPI) with 33 G beveled needles were used to intracranially infuse AAVs into the medial thalamus. The following coordinates were used from Bregma, based on the Paxinos mouse brain atlas, to target these regions of interest: CL thalamus (AP: −1.34mm, ML: −0.75mm, DV: −3.4mm); MDl thalamus (AP: −1.6mm, ML: −0.65mm, DV: −3.6mm). Mice were given a 3-8 week recovery period to allow time for viral spread and transfection to occur. For all surgical procedures in mice, meloxicam (5 mg/kg) was administered subcutaneously at the end of the surgery. All mice were monitored and given meloxicam for up to three days following surgical procedures.

### Spared nerve injury (chronic neuropathic pain model)

As described previously^48^, to induce a chronic pain state in mice, we used a modified version of the Spared Nerve Injury (SNI) model of neuropathic pain. This model involves surgical transection of two branches of the sciatic nerve (common peroneal and tibial branches) while sparing the third (sural branch). To conduct this peripheral nerve injury, anesthesia was induced and maintained throughout a surgery with isoflurane (5% induction, 2% maintenance in oxygen). Each mouse’s left hind leg was shaved and cleaned with ethanol and betadine. We made a 1 cm incision of the skin in the mid-dorsal thigh, approximately where the sciatic nerve branches. The biceps femoris and semimembranosus muscles were gently separated from one another with blunt scissors, which created an opening that exposes the three branches of the sciatic nerve. After that, both the common perotineal and tibial nerves were transected, without suturing with care to not distend the sural nerve. Leg muscles were left uncultured and the skin was closed with a tissue adhesive (3M Vetbond) followed by a Betadine application. During recovery from surgery, mice were placed alone in a cage under a heat lamp until they were awake and ambulating in a normal and balanced manner. Mice were then returned to home cage and closely monitored for three days for well-being.

### Quantification and statistical analysis

All data reported as mean values +/− the standard error of the mean (SEM), unless otherwise stated. Statistical analysis was performed using GraphPad Prism or Python. All data was tested for normality using the Kolgomorov-Smirnoff test. The means of different data distributions were compared using One-Way ANOVA, 2-way ANOVA, Bonferonni’s multiple comparisons, mixed effects model with Bonferonni multiple comparisons, and Tukey’s multiple comparisons. Statistics details for supplemental figures are detailed in supplemental figure legends.

### SHIELD (whole brain clearing, imaging, and analysis)

∼8-week old male and female TRAP2;Sun1-sfGFP double transgenic mice (homozygous for both alleles) were utilized for wholebrain mapping of nociceptive-tagged cells and FOS immunolabeling. Brain samples from all groups were collected and processed on the same day to limit possible batch effects. First, all mice underwent the acute painTRAP protocol to permanently label nociceptive cells with Sun1-sfGFP (a nuclear-envelop protein fused with green fluorescent protein, which was selected to improve automated cell counting by avoiding fluorophores filling cellular processes and obscuring cell somas. Then, 3-days later, mice in the neuropathic-pain groups underwent the SNI neuropathic pain induction procedure for transection of the tibial and common peroneal branches of the left hind limb’s sciatic nerve, while the no-injury groups remained uninjured. Next, 20-days later all mice, in all conditions/groups, were handled and habituated to a testing apparatus for one session. Then at 21-days post-SNI, the nerve-injured mice were returned to the testing apparatus and underwent either 1) chronic pain + light touchFOS: a light-touch mechanical stimulation (0.7g Von Frey hair) protocol to elicit expression of the immediate early gene cFOS consisting of a 1-sec static touch to the lateral edge of the hypersensitivity left hind paw [the spared-sural innervation area], once every 30-sec for 10-min; or 2) chronic painFOS: no stimulation. The no-injury group mice were underwent either 1) acute painFOS: drops of noxious 55°C water (∼50-μl) to the left hind paw, once every 30-sec for 10-min; or 2) controlFOS: no stimulation. After 90-min from the end of the stimulation protocols, all mice were perfused with 10% formalin and 1X PBS and the brains were harvested and fixed overnight in 10% formalin for 24-hr before being transferred to 1X PBS + 0.4% azide. Whole-brain clearing, imaging and automated analysis were performed by LifeCanvas Technologies through a contracted service. In brief, fixed whole brains were prepared with SHIELD (stabilization under harsh conditions via intramolecular epoxide linkages to prevent degradation) to preserve protein antigenicity. Then, the brains were cleared for 7 days with Clear+ delipidation buffer before being immunolabeled with propidium iodide (cell registration), a Goat anti-GFP antibody and a Rabbit anti-FOS antibody using SmartBatch+. Labeled brains were indexed by EasyIndex and imaged by volumetric lightsheet microscopy (SmartSPIM) at 4 uM z-step and 1.8 uM xy pixel size, followed by image post-processing, cell quantification, atlas registration (Allen Institute CCF_v3) and regional graphics. The number and density of FOS^+^, GFP^+^, and the co-expression of FOS^+^ + GFP^+^ cells were calculated for each subregion of the Allen Brain Atlas and visualized as the raw densities or the log2 fold change in density for each medial thalamic subnucleus, relative to the control condition (uninjured; home cage).

### Fluorescent in situ hybridization (RNAScope)

Mice were anesthetized using isoflurane gas in oxygen (5% dose), and the brains were quickly removed and fresh frozen in O.C.T. using Super Friendly Freeze-It Spray (Thermo Fisher Scientific). Brains were stored at −80° C until cut on a Leica cryostat to produce 16 μm coronal sections of the medial thalamus. Sections were adhered to Superfrost Plus microscope slides, and immediately refrozen storing at −80° C. Following the manufacturer’s protocol for fresh frozen tissue for the V2 4Plex RNAscope manual assay (Advanced Cell Diagnostics), slides were fixed for 15-30 min in ice-cold 10% NBF and then dehydrated in a series of ethanol dilutions (50%, 70%, and 100%). Slides were briefly air-dried, and then a hydrophobic barrier was drawn around the tissue sections using a Pap Pen (Vector Labs). Slides were then incubated with hydrogen peroxide solution for 10 min, washed in distilled water, and then treated with the Protease IV solution for 30 min at room temperature in a humidified chamber. Following protease treatment, C1 and C2 cDNA probe mixtures specific for mouse tissue were prepared at a dilution of 50:1, respectively, using the following probes from Advanced Cell Diagnostics: *Oprm1* (C1, 315841), *dsRed* (C2, 481361-C2). Sections were incubated with cDNA probes (2 h at 40°C), and then underwent a series of signal amplification steps using FL v2 Amp 1 (30 min), FL v2 Amp 2 (30 min) and FL v2 Amp 3 (15 min). 2 min of washing in 1x RNAscope wash buffer was performed between each step, and all incubation steps with probes and amplification reagents were performed using a HybEZ oven (ACD Bio) at 40°C. Sections then underwent fluorophore staining via treatment with a serious of TSA Plus HRP solutions and VIVID 570 (1:4000, 7537) and VIVD 650 (1:3000, 7536) dyes from Biotechne TOCRIS. All HRP solutions (C1-C2) were applied for 15 min and Opal dyes for 30 min at 40° C, with an additional HRP blocker solution added between each iteration of this process (15 min at 40° C) and rinsing of sections between all steps with the wash buffer. Lastly, sections were stained for DAPI using the reagent provided by the Fluorescent Multiplex Kit. Following DAPI staining, sections were mounted, and cover slipped using ProLong Gold antifade reagent mounting medium (Invitrogen by Thermo Fischer, P36930, 2936385) and 24×30mm glass coverslips (WVR, 48404-467) before being left to dry overnight in a dark, cool place. Each ISH run included 1 section from all mice with a probe for bacterial mRNA (dapB, ACD Bio, 310043) to serve as a negative control.

### Imaging and Quantification

All tissue was imaged on a Keyence BX-X810 all-in-one fluorescence slide-scanning microscope at 48-bit resolution using the following objectives: PlanApo_<monospace>λ </monospace>x4 0.20/20.00mm, PlanApo_<monospace>λ </monospace>x20 0.75/1.00mm, and PlanApo_<monospace>λ </monospace>60xH 1.40/0.13mm oil objective. All imaging processing and stitching prior to quantification was performed with the Keyence BZ-X analyzer software (version 1.4.0.1). For cell body quantification, every third slice across the whole brain was collected, amplified with IHC, and imaged at 4X for initial visualization. High-density areas were imaged at 20X with Z-stacks. For each ROI, multiple slice coordinates were selected per mouse that spanned the anterior-posterior boundaries of the ROI. Quantification of neurons expressing fluorophores was performed either via manual counting of TIFF images in Photoshop (Adobe, 2021) using the Counter function or ImageJ counter function, or via HALO AI Cell counting software. For axon density quantification, immunohistochemistry was performed to amplify the mCherry signal and visualize MTh^MOR^ axons throughout the brain in 30 uM tissue free-floating slices. Areas with dense axon innervation were identified using 4X imaging. Areas with most dense axon innervation were selected for additional 20X imaging with Z-stacks. The exposures for FITC and CY3 were adjusted to avoid overexposed pixels for the brightest area. This exposure was kept consistent for all slices for an individual mouse. For an individual ROI, three slices per mouse were included. ROIs were drawn in Adobe Illustrator and axon density was quantified with AxoDen^65^.

## Supplemental Figures

**Fig. S1.**
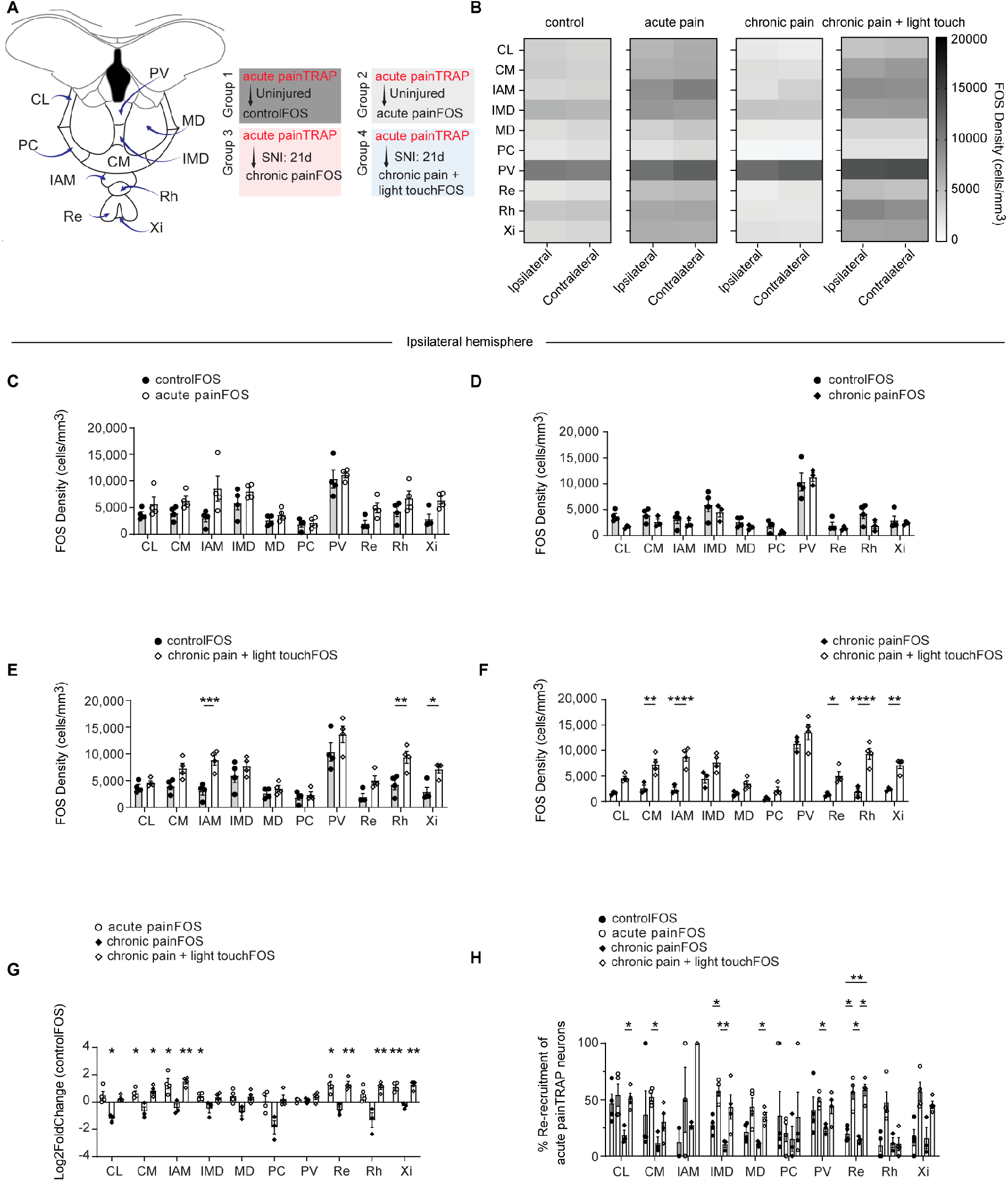
Acute nociception and allodynia increase ipsilateral MTh activity. (A) (Left) Coronal cross-section atlas of medial thalamus annotated with the ten MTh subnuclei, each split into ipsilateral (same hemisphere as stimulus) and contralateral (opposite hemisphere as stimulus) parts. (Right) Breakdown of four experimental FOS stimulation groups: controlFOS= no nerve injury + home cage conditions. acute painFOS= no nerve injury, noxious hot water drop to left hind paw stimulation. chronic painFOS: nerve injury + home cage conditions. chronic pain + light touchFOS: nerve injury + 0.7g von Frey stimulation to left hind paw. All mice received acute painTRAP. (B) Heat maps showing FOS density in all four groups in each MTh nucleus on the ipsilateral and contralateral hemispheres. (C) FOS density comparing controlFOS with acute painFOS conditions. (D) FOS density comparing controlFOS with chronic painFOS groups. (E) FOS density comparing controlFOS with chronic pain + light touchFOS groups. (F) FOS density comparing chronic painFOS and chronic pain + light touchFOS groups. (G) Log2Fold change in FOS density for each group compared to the controlFOS condition. One sample t-test vs. zero determined significance. *p<0.05, **p<0.01. (H) Percent of acute painTRAP^sf-GFP^ labeled cells that co-label with FOS in each group. (B-F; H) Two-way ANOVA with Bonferonni multiple comparisons determined significance. *p<0.05, **p<0.01, ***p<0.001 (B-H) Bars represent the mean +/− SEM for n=3-4 TRAP2:CagSun1^sfGFP^ per group; individual data points represent each mouse. Abbreviations: CL, central lateral. CM, central medial nucleus. IAM, interanteromedial nucleus. IMD, intermediodorsal nucleus. MD, mediodorsal thalamus PC, paracentral nucleus. PV, paraventricular nucleus. Re, nucleus reuniens. Rh, rhomboid nucleus. Xi, xiphoid nucleus.

**Fig. S2.**
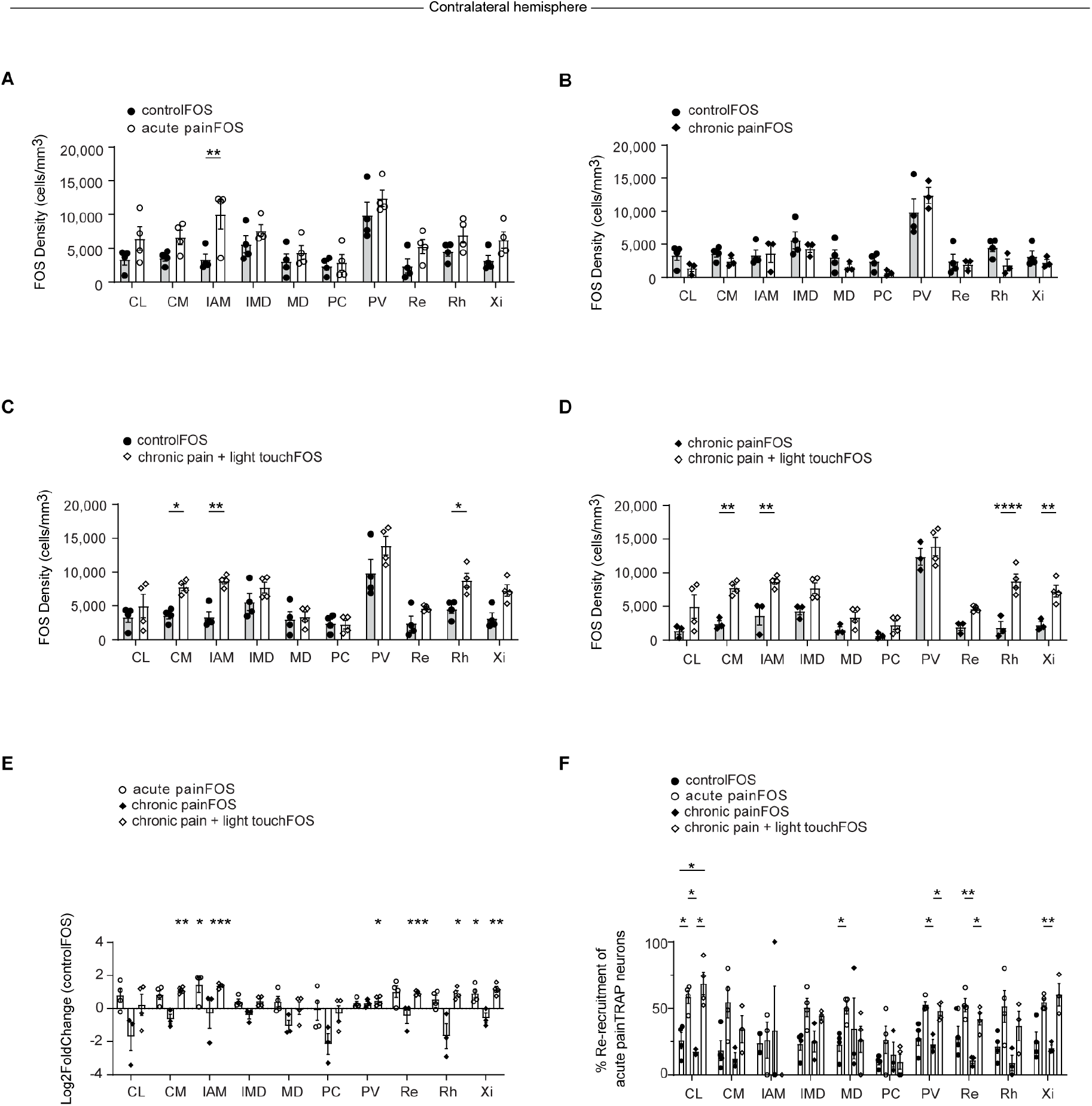
Acute nociception and allodynia increase contralateral MTh activity. (A) FOS density comparing controlFOS with acute painFOS conditions. (B) FOS density comparing controlFOS with chronic painFOS groups. (C) FOS density comparing controlFOS with chronic pain + light touchFOS groups. (D) FOS density comparing chronic painFOS and chronic pain + light touchFOS groups. (E) Log2Fold change in FOS density for each group compared to the controlFOS condition. One sample t-test vs. zero determined significance. *p<0.05, **p<0.01. ***p<0.001. (F) Percent of acute painTRAP^sf-GFP^ labeled cells that co-label with FOS in each group, calculated using: (number of co-labeled cells / number of painTRAP^sf-GFP^ cells)*100%. (A-D; F) Two-way ANOVA with Bonferonni multiple comparisons determined significance. *p<0.05, **p<0.01, ***p<0.001 (A-F) Bars represent the mean +/− SEM for n=3-4 TRAP2:CagSun1^sfGFP^ per group; individual data points represent each mouse. Abbreviations: CL, central lateral. CM, central medial nucleus. IAM, interanteromedial nucleus. IMD, intermediodorsal nucleus. MD, mediodorsal thalamus PC, paracentral nucleus. PV, paraventricular nucleus. Re, nucleus reuniens. Rh, rhomboid nucleus. Xi, xiphoid nucleus. Contralateral hemisphere= opposite hemisphere of stimulus.

**Fig. S3.**
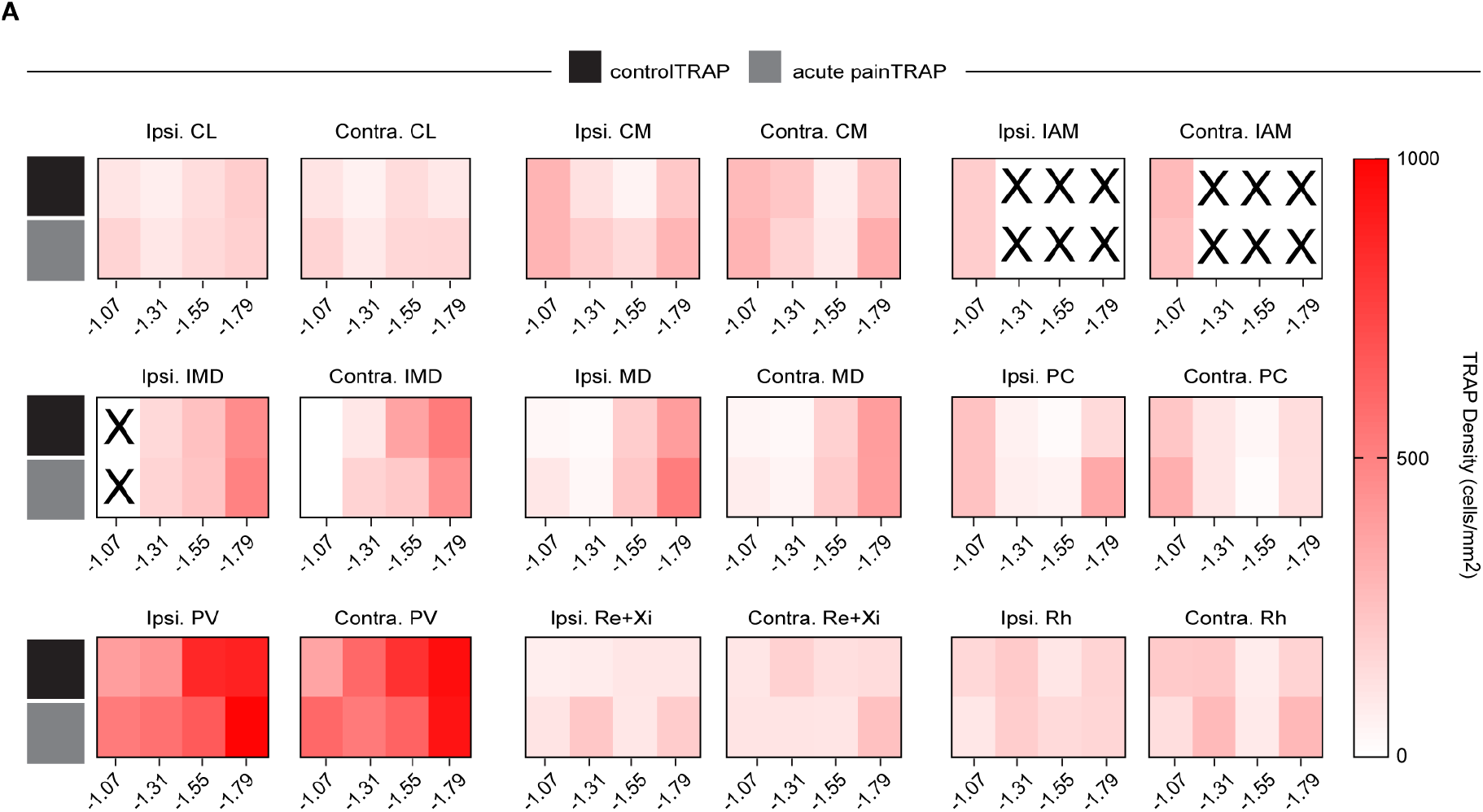
No effect of acute pain found in anterior-posterior analysis of acute painTRAP vs. controlTRAP density. (A) Heat maps for mean TRAP2^Ai9^ densities in controlTRAP and acute painTRAP groups at −1.07, −1.31, −1.55, −1.79mm from Bregma in each MTh subnucleus. Independent t-tests between controlTRAP and acute painTRAP at each coordinate determined significance for acute painTRAP^Ai9^ (n=11 females, n=5 males) and controlTRAP^Ai9^ (n=11 females, n=4 males) mice. Ipsilateral= same hemisphere as acute pain stimulus. Contralateral= opposite hemisphere as acute pain stimulus. Abbreviations: CL, central lateral. CM, central medial nucleus. IAM, interanteromedial nucleus. IMD, intermediodorsal nucleus. MD, mediodorsal thalamus PC, paracentral nucleus. PV, paraventricular nucleus. Re+Xi, averaged across nucleus reuniens and xiphoid nucleus. Rh, rhomboid nucleus.

**Fig. S4.**
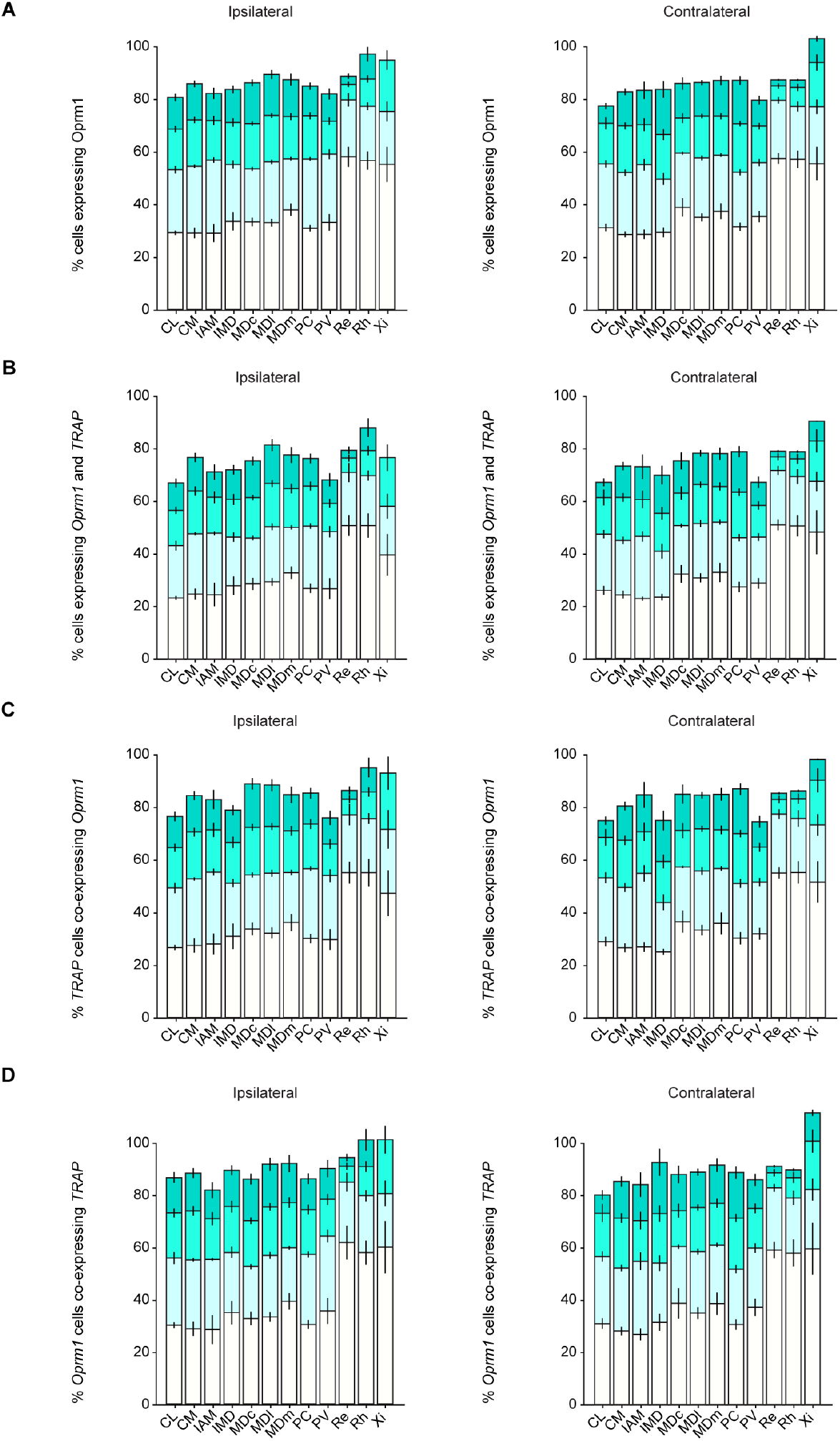
Transcriptomic analysis of *Oprm1*^*+*^ and acute pain*TRAP*^*Ai9+*^ populations in MTh are not lateralized. (A) Quantification of the percent of cells expressing *Oprm1* in each MTh subnucleus (Number of *Oprm1*^+^ cells/total number of cells)*100%. (B) Quantification of the percent of cells expressing both *Oprm1* and *acute painTRAP*^*Ai9*^ in each MTh subnucleus (Number of *acute painTRAP*^*Ai9*^ and *Oprm1* co-expressing cells/total number of cells)*100%. (C) Quantification of the percent of *acute painTRAP*^*Ai9+*^ cells co-expressing *Oprm1* (Number of *acute painTRAP*^*Ai9*^ and *Oprm1* co-expressing cells/total number of TRAP^*Ai9*+^ cells)*100%. (D) Quantification of the percent of *Oprm1*^+^ cells co-expressing *acute painTRAP*^*Ai9*^ (Number of *acute painTRAP*^*Ai9*^ and *Oprm1* co-expressing cells/total number of *Oprm1*^+^ cells)*100%. (A-D) (Left graph= hemisphere ipsilateral to acute painTRAP stimulus. Right graph= hemisphere contralateral to acute painTRAP stimulus. Bars represent the mean +/− SEM for n=3 male and n=3 female TRAP2:Ai9 mice. Abbreviations: CL, central lateral. CM, central medial nucleus. IAM, interanteromedial nucleus. IMD, intermediodorsal nucleus. MDc, mediodorsal nucleus, central part. MDl, mediodorsal nucleus, lateral part. MDm, mediodorsal nucleus, medial part. PC, paracentral nucleus. PV, paraventricular nucleus. Re, nucleus reuniens. Rh, rhomboid nucleus. Xi, xiphoid nucleus.

**Fig. S5.**
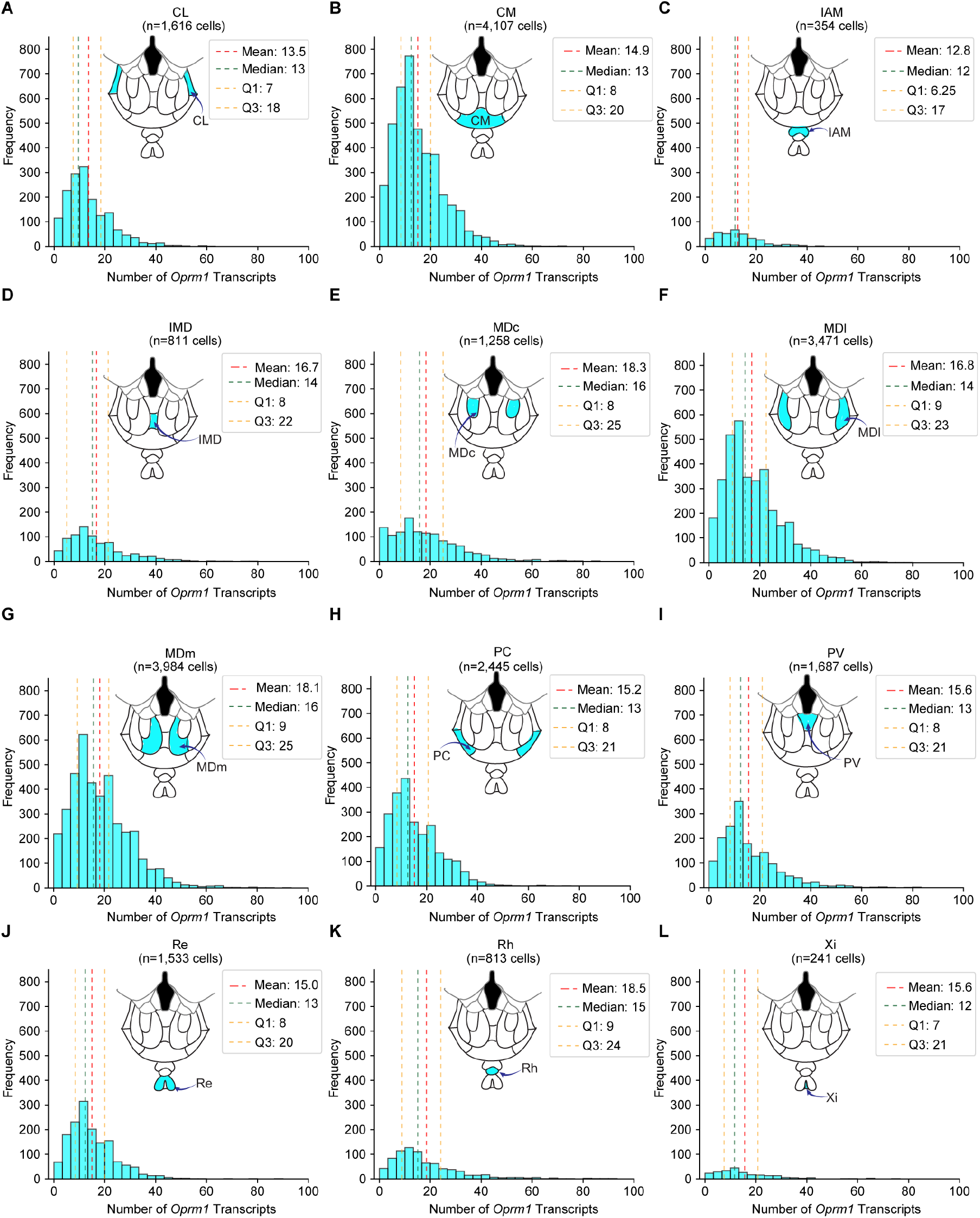
Heterogeneous distribution of *Oprm1* transcripts in each MTh subnucleus. (A-L) Histogram for Number of *Oprm1* transcripts in each MTh subnucleus, summed across both hemispheres of n=3 female and n=3 male TRAP2:Ai9 mice. Abbreviations: CL, central lateral. CM, central medial nucleus. IAM, interanteromedial nucleus. IMD, intermediodorsal nucleus. MDc, mediodorsal nucleus, central part. MDl, mediodorsal nucleus, lateral part. MDm, mediodorsal nucleus, medial part. PC, paracentral nucleus. PV, paraventricular nucleus. Re, nucleus reuniens. Rh, rhomboid nucleus. Xi, xiphoid nucleus.

**Fig. S6.**
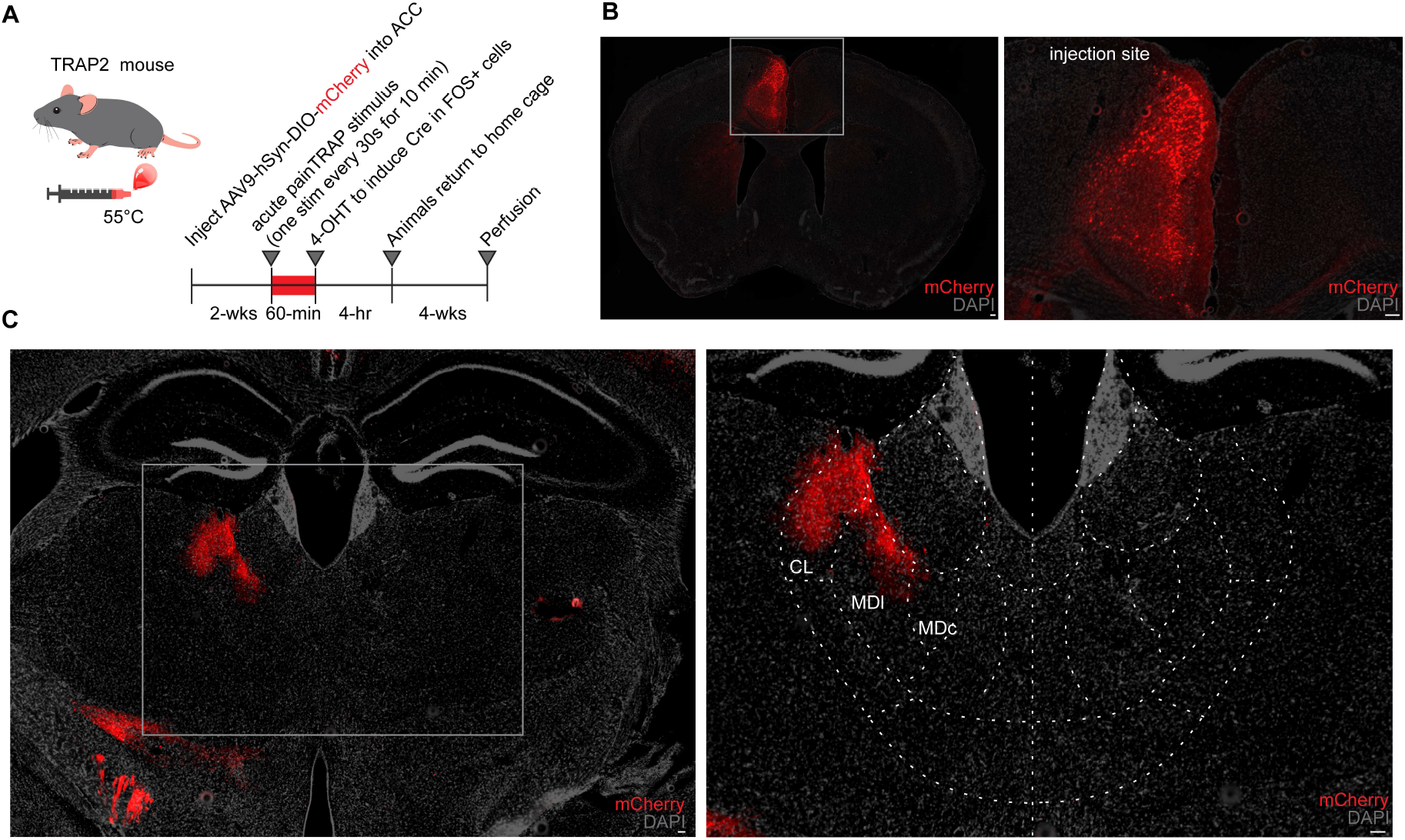
Nociceptive neurons in the anterior cingulate cortex send axons to CL and MD thalamus. (A) Experimental design for *acute painTRAP* protocol with the injection of a cre-dependent mCherry virus to label nociceptive-active ACC neurons in *TRAP*2 mouse. (B) 20x representative image of ACC injection site. (Left) whole-slice image. (Right) ACC-zoomed image. (C) 20x representative image of viral expression in thalamus. (Left) whole-slice image. (Right) MTh-zoomed image. Abbreviations: ACC, anterior cingulate cortex, CL, central lateral nucleus, MDl, mediodorsal nucleus, lateral part. MDc, mediodorsal nucleus, central part. MTh= medial thalamus. Scale bars: 250 µM.

**Fig. S7.**
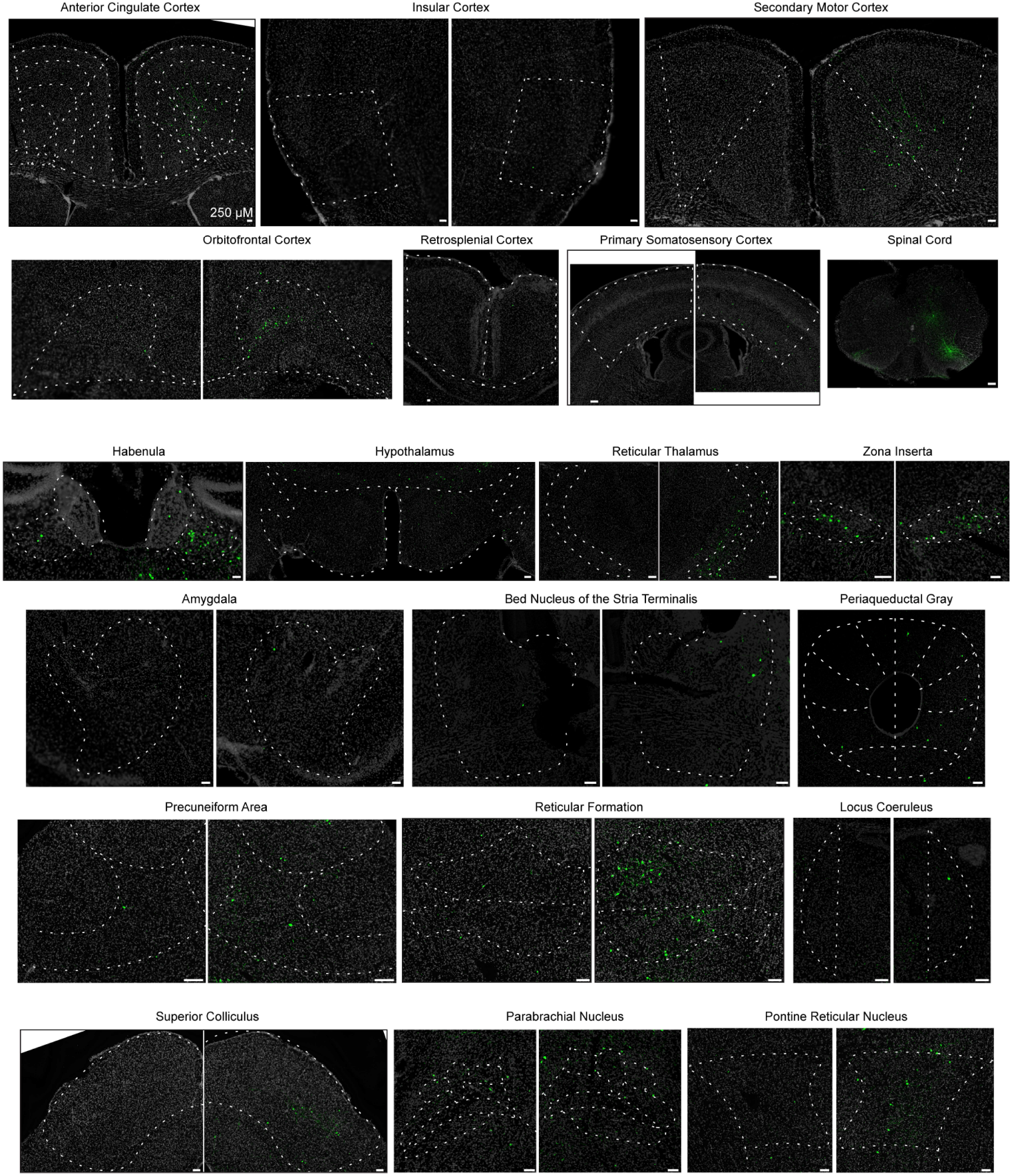
Bilateral brain-wide monosynaptic inputs to MTh^MOR^. 20x representative inputs of brain-wide, bilateral inputs to MTh^MOR^ neurons. Input neurons are labeled in green from injection of EnvA-pseudotyped, G-deleted, GFP-expressing rabies virus (RabV-GFP) into MTh. Images sourced from slices collected from n=4 female, n=2 male *Oprm1*-T2A-Cre mice. Scale bars: 250 µM.

**Fig. S8.**
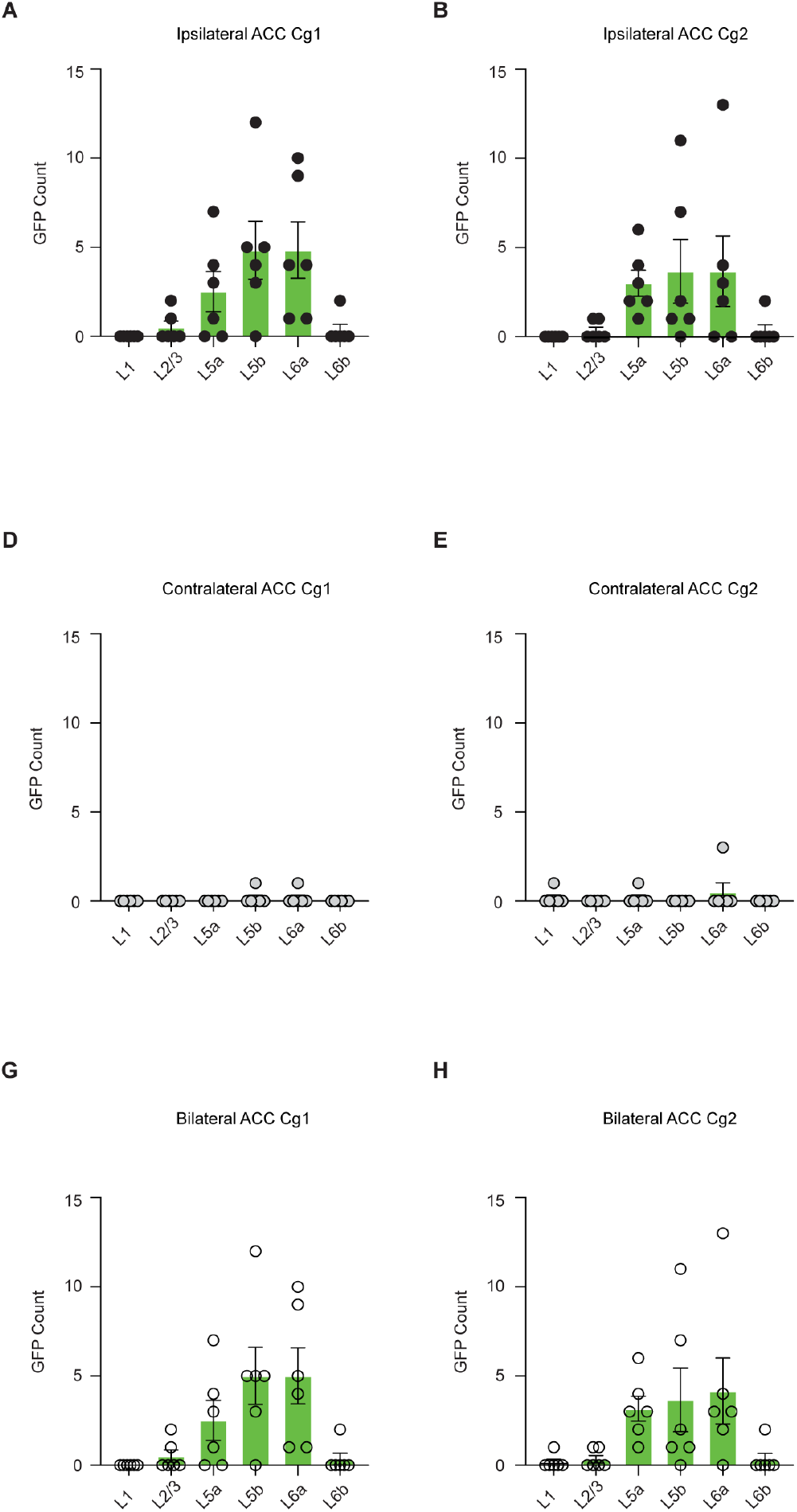
No subregion-specific differences in inputs to MTh^MOR^ from ACC. (A) Comparison of GFP count in each layer of ACC Cg1 ipsilateral to viral injection. (B) Comparison of GFP count in each layer of ACC Cg2 ipsilateral to viral injection.(C) Comparison of GFP count in each layer of ACC Cg1 contralateral to viral injection. (D) Comparison of GFP count in each layer of ACC Cg2 contralateral to viral injection. (E) Comparison of GFP count in each layer of ACC Cg1, calculated by averaging left and right hemisphere counts. Comparison of GFP count in each layer of ACC Cg2, calculated by averaging left and right hemisphere counts. (A-E) Bars represent the mean +/− SEM; data points for individual n=4 female and n=2 male *Oprm1*-T2A-Cre mice. One-way ANOVA with Bonferonni multiple comparisons determined significance. Abbreviations: ACC, anterior cingulate cortex (Cg1, prelimbic area. Cg2, infralimbic area). L1, cortical layer 1. L2/3, cortical layers 2/3. L5a, cortical layer 5a. L5b, cortical layer 5b. L6a, cortical layer 6a. L6b, cortical layer 6b).

**Fig. S9.**
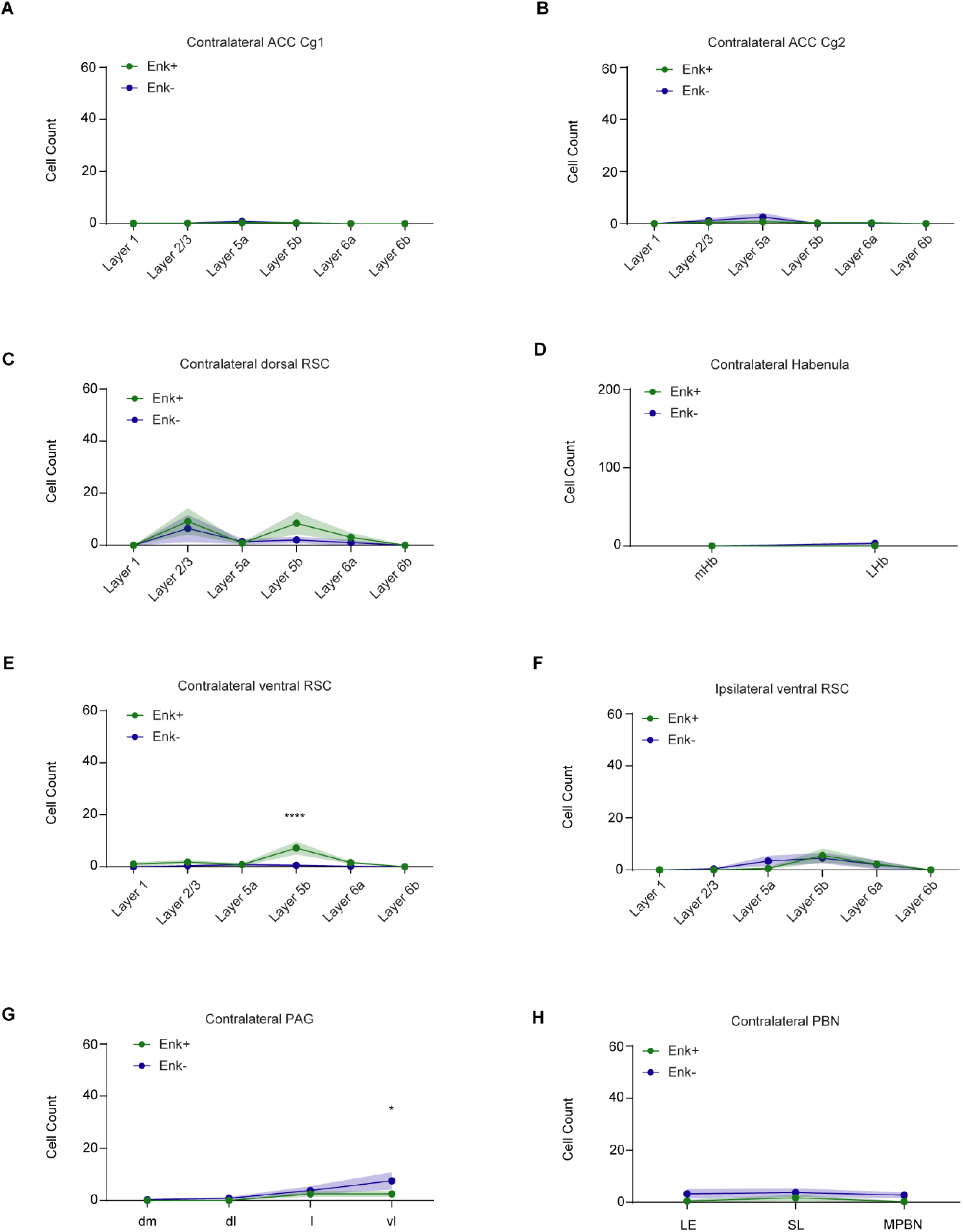
RSC and PAG show subregion differences of Enk+ vs. Enk− inputs to MTh. (A-H) Quantification of raw number of GFP-expressing and mCherry-expressing cells in subregions of ACC, PAG, PBN, Habenula, and RSC. Two-way ANOVA with Bonferonni multiple comparisons determined significance (*p<0.05, **p<0.01, ***p<0.001 ****p<0.0001) for n=6 male Penk-Cre mice. Ipsilateral=hemisphere on the same side as acute painFOS stimulus. Contralateral=hemisphere on the opposite side of acute painFOS stimulus. Points represent mean +/− SEM for n=6 male *Penk*-Cre mice. Abbreviations: ACC, anterior cingulate cortex (Cg1, prelimbic area. Cg2, infralimbic area). Habenula (LHb, lateral habenula. mHb, medial habenula). PAG, periaqueductal gray (dm, dorsomedial column. dl, dorsolateral column. l, lateral column. vl, ventrolateral column). PBN, parabrachial nucleus (LE, external lateral nucleus. SL, superior lateral nucleus. MPBN, medial parabrachial nucleus).

**Fig. S10.**
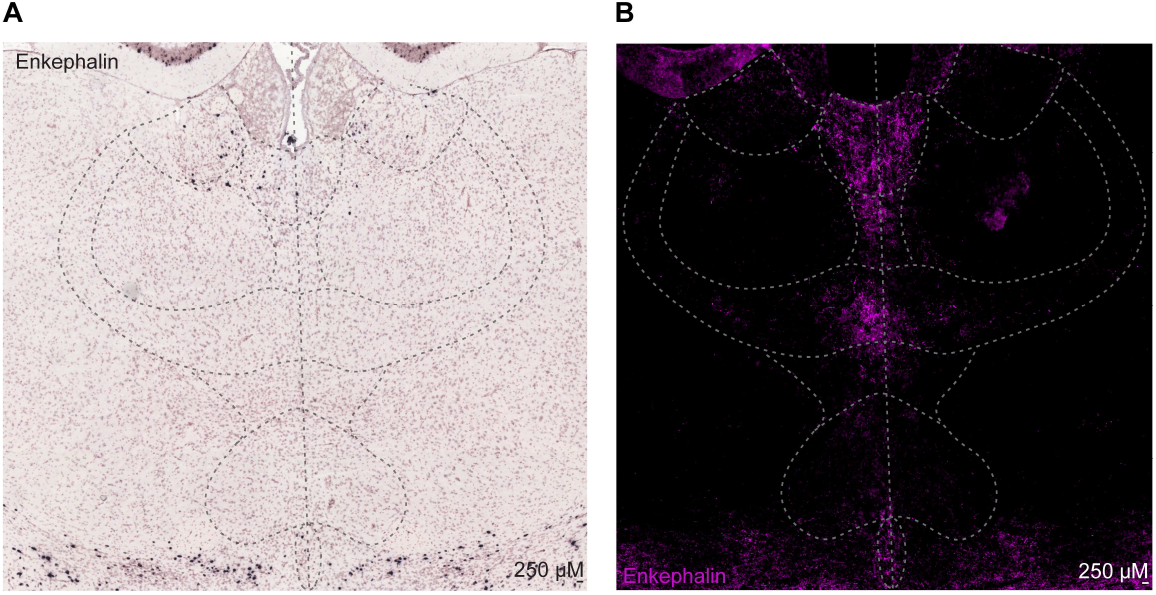
Expression of enkephalin in axons within MTh, not MTh somas. (A) Representative image of pro-enkephalin mRNA within MTh. Image acquired from Allen Mouse Brain Atlas *In situ hybdridization* data. (B) 20x representative image of enkephalin protein expression within MTh. Image sourced from slices collected from n=3 male C57BL6/J mice. Scale bars: 250 µM.

